# Acquisition and loss of secondary metabolite clusters shaped the evolutionary path of three recently emerged phytopathogens of wheat

**DOI:** 10.1101/283416

**Authors:** Elisha Thynne, Oliver L. Mead, Yit-Heng Chooi, Megan C. McDonald, Peter S. Solomon

## Abstract

- White grain disorder is a recently emerged wheat disease in Australia, caused by three *Botryosphaeriaceae spp.*; *Eutiarosporella darliae, E. pseudodarliae*, and *E. tritici-australis*. The disease cycle of these pathogens and the molecular basis of their interaction with wheat are poorly understood. To address this, we undertook a comparative genomics approach to identify potential pathogenicity factors.
- Subsequent genome analysis revealed that each of the white grain disorder species harbour modular polyketide synthase genes. To our knowledge, this is the first report of fungi harbouring such genes. Further comparative analysis using the modular polyketide synthase genes discovered their presence in the closely related *Macrophomina phaseolina*. Phylogenetic analysis implicates horizontal acquisition of these genes from a bacterial or a protist species.
- Both *E. darliae* and *E. pseudodarliae* possess a secondary metabolite cluster with multiple polyketide/non-ribosomal peptide synthase genes (*Hybrid-1, -2, and -3*). In contrast, only remnant and partial genes homologous to this cluster were identified at a syntenic locus in *E. tritici-australis* suggesting loss of this cluster. Homologues of *Hybrid-2* in other fungi have been proposed to facilitate disease induction in woody plants. Subsequent assays confirmed that *E. darliae* and *E. pseudodarliae* were both pathogenic on woody plant hosts, but *E. tritici-australis* was not, implicating woody plants as potential host reservoirs for the fungi. We hypothesise that loss of the cluster in *E. tritici-australis* represents a committed lifestyle jump to grasses.
- Combined, our observations relating to the secondary metabolite potential of the WGD *Eutiarosporella spp.* have contributed novel data to the field by expanding the range of known fungal secondary metabolite genes, and helped develop our understanding of the lifestyle and potential host-range of a recently emerged pathogen of wheat.

## Introduction

Fungal species occupy a wide range of trophic niches across the world. Whilst many blend into the environment unseen, a small but significant number are voracious phytopathogens, responsible for substantial losses to agricultural, horticultural, and forestry industries (Agrios, 2005). Understanding how fungal species evolved to persist in the hostile environment of a host plant, whether as an endophyte or a pathogen, is a major focus of mycological research.

The Dothideomycetes are a prominent class of fungi encompassing many economically significant phytopathogens. In recent years, a large number of Dothideomycete species’ genomes have been sequenced and analysed (Ohm *et al*., 2012). Further, the complex mechanisms that these fungi use to facilitate growth in a host plant and induce disease are being teased apart (Oliver and Solomon, 2010). Small molecules, known as secondary metabolites (SMs) are often used to enable unique trophic lifestyles (Friesen *et al*., 2008, Walton, 1996, Stergiopoulos *et al*., 2013). These small molecules can promote growth in environments where there is strong competition from other microbes or environmental stress (Shabuer *et al*., 2015, Stergiopoulos *et al*., 2013). The Dothideomycetes produce a wide range of SMs (Muria - Gonzalez *et al*., 2015), some of which are deployed as host specific toxins (HSTs) that facilitate virulence in a genotype specific manner (Friesen *et al*., 2008, Walton, 1996, Stergiopoulos *et al*., 2013). A selection of Dothideomycete SM HSTs include; HC-toxin, a non-ribosomal peptide synthase (NRPS) produced by *Cochliobolus carbonum* (Walton, 2006); T-toxin, a polyketide synthase (PKS) produced by *Cochliobolus heterostrophus* (Yang *et al*., 1996, Baker *et al*., 2006); and ACT toxin and AK toxin, two PKSs produced by tangerine and Japanese pear pathovars of *Alternaria alternata*, respectively (Nakashima *et al*., 1982, Kohmoto *et al*., 1993, Tanaka *et al*., 1999, Miyamoto *et al*., 2010).

White grain disorder (WGD) is a disease of wheat first observed in Queensland, Australia during the late 1990s causing the shrivelling and discoloration of grain-heads (Wildermuth *et al*., 2001, Platz, 2011). In the past decade, the prevalence of WGD has spread within Australia and the disease has now been observed in wheat-growing regions in Western Australia, South Australia, and Queensland (Thomas and Jayasena, 2015, Evans, 2013). The causal agents of WGD are three closely related species; *Eutiarosporella darliae*, *E. pseudodarliae*, and *E. tritici-australis* (Thynne *et al*., 2015). These are Dothideomycete fungi in the Botryosphaeriaceae family. It is unknown how these three species emerged as wheat pathogens, where they originated from, and if they interact with any non-grass hosts.

In this study, we have undertaken a comparative genomics analysis of the three WGD *Eutiarosporella spp* to gain an insight into their evolution and emergence as wheat pathogens. Many Dothideomycete genomes are now sequenced and available for comparative purposes (Ohm *et al*, 2012). A shared theme among the study of these genomes is that genomic adaptation events specialise these fungi into their respective lifestyle niches (Friesen *et al*, 2006, Goodwin, *et al*, 2011, Ohm *et al*, 2012, Stuckenbrock *et al*, 2012, Dhillon *et al*, 2015). As such, we hypothesised that genomic adaptation events played a role in specialising the WGD disorder species to explain their current lifestyle. We focused our analyses upon genomic adaptation events that have shaped the SM capabilities of the WGD *Eutiarosporella spp.* and apply knowledge about these genes to extrapolate potential trophic niches.

## Results

### The WGD *Eutiarosporella spp*. have the smallest assembled Dothideomycete genomes, to date, and have reduced secondary metabolite potential

The *de novo* genome assemblies of *E. darliae*, *E. pseudodarliae*, and *E. tritici-australis* used in this analysis were approximately 27 Mb and predicted to encode between 8500 and 8750 genes (Table 1). The genomes of the three WGD species are reduced in size and gene content in comparison with the genomes of other sequenced members of the Botryosphaeriaceae, and other described members of the Dothideomycetes. Comparisons against the genome assemblies of the Botryosphaeriaceae species; *Botryosphaeria dothidea, Diplodia seriata, Neofussicoccum parvum*, and *Macrophomina phaseolina* are displayed in *Table 1* (Blanco-Ulate *et al*., 2013, Islam *et al*., 2012, van der Nest *et al*., 2014, Morales-Cruz *et al*., 2015). We also include three other Dothideomycete species; *Zymoseptoria tritici*, *Mycosphaerella populorum*, and *Mycosphaerella populicola* for additional comparison with species outside of the Botryosphaeriaceae. (Dhillon *et al*., 2015, Ohm *et al*., 2012). The latter two species are putatively the next smallest Dothideomycete genomes assembled to date (Dhillon *et al*, 2015). The Core Eukaryotic Gene Mapping Approach (CEGMA) scores for estimating genome assembly completeness are high for each of the *<u>Eutiarosporella</u>* <u>spp.</u> genomes (>96%) (Table 1). This suggests that, although the genome assemblies are smaller than those of the other Dothideomycete species listed, these genome assemblies are relatively complete (Parra *et al*., 2007).

**Table 1:**
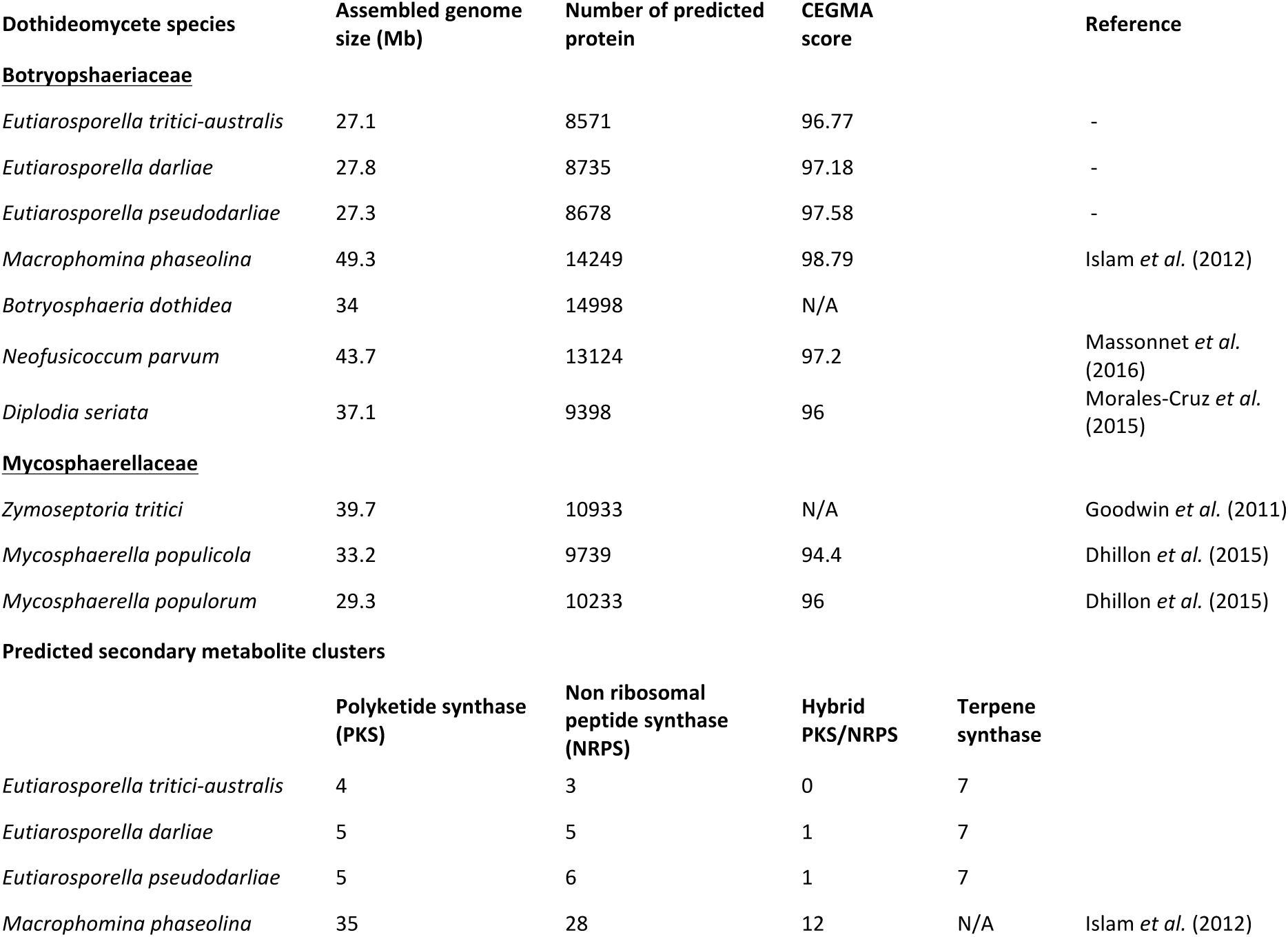
Basic genome statistics.

The predicted protein sets for each of the WGD species were submitted to the antiSMASH online server (v2.0) for secondary metabolite prediction (Weber *et al*., 2015a). The predicted SM clusters are listed in Table 1 and the specific proteins in Table 2. For comparative purposes, we have included *M. phaseolina’s* predicted SM gene counts (Islam *et al*., 2012) (Table 1). Similar to overall predicted gene numbers, the *Eutiarosporella* spp. have a reduced SM complement to *M. phaseolina*. BLASTp was used to compare the SM sequences against the NCBI and JGI databases in order to determine the distribution of homologues among other organisms (Table 2). From this list, most of the secondary metabolite amino acid sequences have best-BLAST hits to sequences from other members of the Botryosphaeriaceae. However, among the NRPSs identified, three had best-BLAST hits from Sordariomycetes: NRPS2 had a best-BLAST hit to *Helicocarpus griseus*; NRPS3, which is absent in *E. tritici-australis*, had a best blast hit from *Trichophyton benhamiae*; NRPS5, which is only present in *E. pseudodarliae*, had a best-BLAST hit to an NRPS from *Colletotrichum gloeosporoides*. However, each of these NRPSs only share low sequence identity with these putative homologues (<50% shared identity). The Terpene synthases (TSs) and iterative PKSs (iPKSs) are largely conserved among the three *Eutiarosporella* species, suggesting that most SM genes are vertically transmitted among these three species. The PKS-NRPS hybrid genes appeared to be exclusively absent in *E. tritici-australis* among the three. The biggest surprise from secondary metabolite gene cluster analysis is that multiple modular PKSs, which are commonly found in bacteria and protists but not fungi, were identified in the three *Eutiarosporella* genomes (Table 1, 2). The origins of these modular PKSs and PKS-NRPSs are discussed further below.

**Table 2.** Secondary metabolite genes.

| Individual SM<br>synthase genes | Presence/absence<br>(+/-) |  |  | Best-BLAST<br>species | Accession | Identity | Predicted<br>SM type |
| --- | --- | --- | --- | --- | --- | --- | --- |
| TSs | *E. tritici-*<br>*australis* | *E.*<br>*darliae* | *E.*<br>*pseudodarliae* |  |  |  |  |
| 1 | + | + | + | *D. seriata* | KKY22841.1 | 86%_identity | TS |
| 2 | + | + | + | *D. corticola* | OJD35063.1 | 70%_identity | TS |
| 3 | + | + | + | *N. parvum* | XP_007581866.1 | 86%_identity | TS |
| 4 | + | + | + | *D. seriata* | OMP85614.1 | 85%_identity | TS |
| 5 | + | + | + | *D. corticola* | XP_020126315.1 | 78%_identity | TS |
| 6 | + | + | + | *D. seriata* | KKY14050.1 | 64%_identity | TS |
| 7 | + | + | + | *N. parvum* | EOD50659.1 | 86%_identity | TS |
| NRPSs |  |  |  |  |  |  |  |
| 1 | + | + | + | *D. corticola* | XP_020135006.1 | 73%_identity | NRPS |
| 2 | + | + | + | *H. griseus* | PGH17632.1 | 37%_identity | NRPS |
| 3 | - | + | + | *T. benhamiae* | DAA73007.1 | 48%_identity | NRPS |
| 4 | - | + | + | *N. parvum* | EOD50138.1 | 66%_identity | PKS-NRPS |
| 5 | - | - | + | *C. gloeosporioides* | EQB52787.1 | 44%_identity | NRPS |
| PKS-NRPSs |  |  |  |  |  |  |  |
| 1 | - | + | + | *M. phaseolina* | EKG15176.1 | 90%_identity | PKS-NRPS |
| 2 | - | + | + | *M. phaseolina* | EKG15173.1 | 85%_identity | PKS-NRPS |
| PKSs |  |  |  |  |  |  |  |
| 1 | + | + | + | *D. corticola* | XP_020132262.1 | 74%_identity | PKS |
| 2 | + | + | + | *D. seriata* | KKY20157.1 | 90%_identity | PKS |
| 3 | + | + | + | *N. parvum* | EOD47055.1 | 80%_identity | PKS |
| mPKSs |  |  |  |  |  |  |  |
| a | + | + | + | *M. phaseolina* | EKG12504.1 | 68%_identity | mPKS |
| b | + | + | + | *M. phaseolina* | EKG12506.1 | 73%_identity | mPKS |
| c | - | + | + | *M. phaseolina* | EKG12504.1 | 74%_identity | mPKS |
| d | - | - | + | *M. phaseolina* | EKG12506.1 | 67%_identity | mPKS |

### The modular PKS genes in the wheat-infecting *Eutiarosporella spp.* exist in pairs with missing acyltransferase domain in some modules

Based on antiSMASH secondary metabolite cluster predictions, four modular PKS genes (*mPKSs*) were identified within the WGD *Eutiarosporella* genomes (named a, b, c, and d) (Fig. 1 a, b, c). mPKSs are typically found in bacteria or protist species, and differ from the typical fungal iPKSs, in that each enzymatic domain can be represented many times in a single protein, forming modules of enzymatic domains. However, each module of domains is only used once during the synthesis of the metabolite product and the ‘growing’ intermediates are passed to subsequent module in an assembly line manner (Robbins et al., 2016). In contrast, iPKSs only harbour one copy of each enzymatic domain, however, these could be used multiple times during the synthesis of a metabolite product in an iterative manner (Chooi et al. 2015). iPKSs are the hall mark of fungal PKSs, although some bacterial PKSs are known to be iterative (Chen and Du 2016).

**Figure 1:**
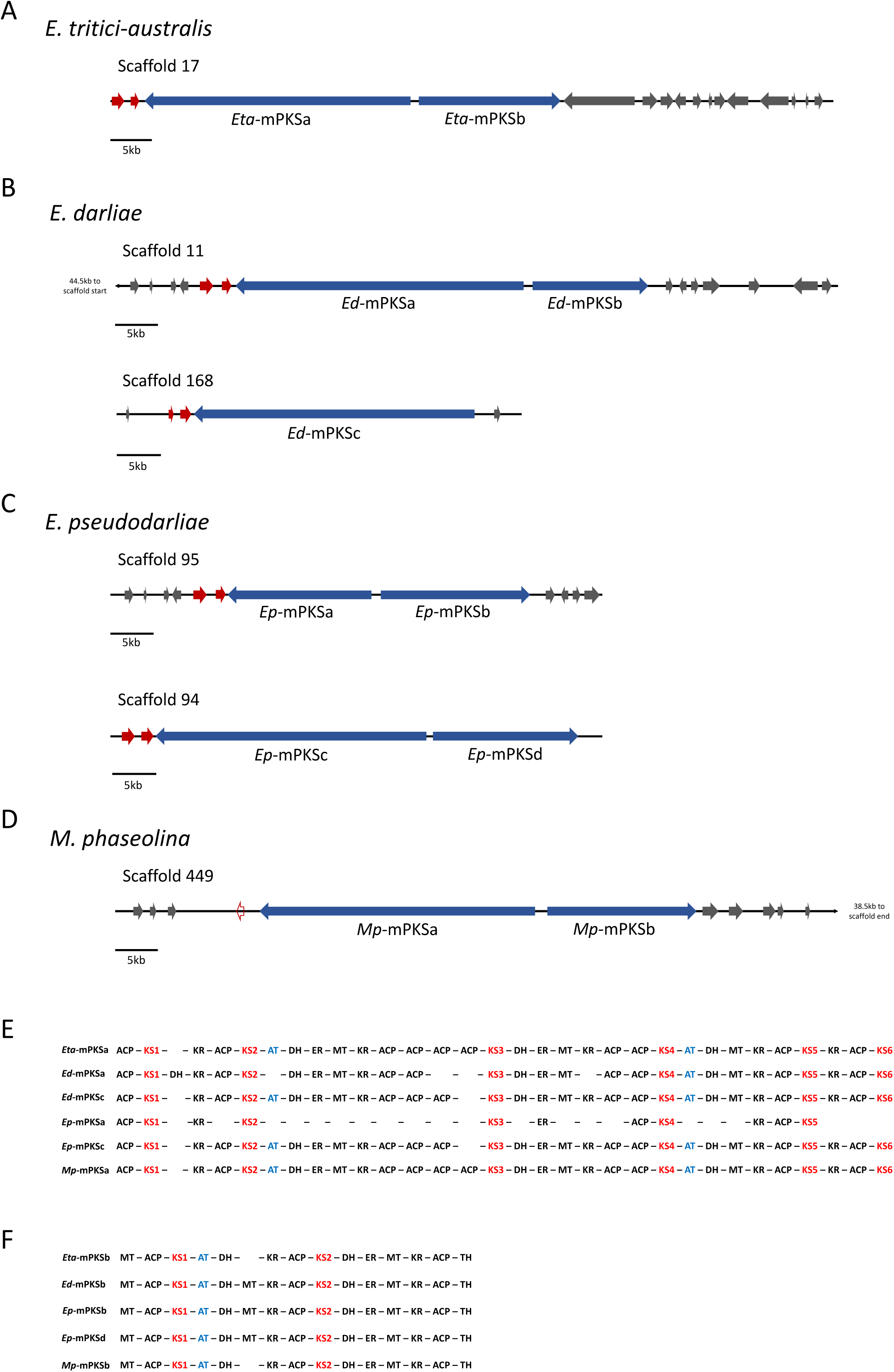
The WGD *Eutiarosporella spp.* and *M. phaseolina* harbour modular PKS genes (mPKS). Abreviations: ACP (acyl carrier protein), KS (ketoacyl synthase), DH (dehydratase), KR (ketoreductase), AT (ACP transacylase), ER (enoyl reductase), MT (methyltransferase), TH (thiohydrolase). KS and AT are each highlighted (red and blue, respectively) as they are discussed in greater detail in the text. A) Graphical representations of *E. tritici-australis* mPKS encoding genes, *Eta-*mPKS-a and *Eta-*mPKS-b. B) Graphical representations of *E. darliae* mPKS encoding genes, *Ed-*mPKS-a, *Ed-*mPKS-b, and *Ed-*mPKS-c. C) Graphical representations of *E. pseudodarliae* mPKS encoding genes, *Ep-*mPKS-a, *Ep-*mPKS-b, *Ep-*mPKS-c, and *Ep-*mPKS-d. D) Graphical representations of *M. phaseolina* mPKS encoding genes, *Mp-*mPKS-a and *Mp-*mPKS-b. E) Alignment of mPKSa and mPKSc protein sequence domains, highlighting comparative presence absence of domains. F) Alignment of mPKSb and mPKSd protein sequence domains, highlighting comparative presence absence of domains. (Note: domain alignments are not based off phylogenetic relationship, but merely the order in which they occur).

We display the gene orientation and encoded protein domains of each of the *Eutiarosporella* mPKSs in *Figure 1*. To describe the structure of the *Eutiarosporella* mPKS genes, we refer to their Ketoacyl synthase (KS) domains. KS domains are responsible for polyketide chain elongation in PKS systems and in mPKSs, they are also involved in intermodular translocation of the growing polyketide chain (Robbins et al., 2016). As such, they are the essential functional domain in PKSs and are present in each module within mPKSs. *E. tritici-australis* has two mPKS genes that share a 5’ non-coding region on opposite strands (*Eta-mPKS*-a and *Eta-mPKS*-b) (Fig. 1a). The encoded amino-acid sequences contain six and two KS domains, respectively (Fig. 1e, f). *E. darliae* has three mPKS genes; a pair, that share a 5’ non-coding region and are predicted on opposite strands (*Ed-mPKSa* and *Ed-mPKS*b) and a solo mPKS gene (*Ed-mPKSc*) (Fig. 1b). The encoded amino acid sequences contain six (a) and two (b), and six (c) KS domains, respectively (Fig. 1e, f). *E. pseudodarliae* has four mPKS genes; two pairs, each that share a 5’ non-coding region and are predicted on opposite strands (*Ep-mPKSa* and *Ep-mPKSb*, and, *Ep-mPKSc* and *Ep-mPKSd*) (Fig. 1c). The first pair of mPKS genes contain five (a) and two (b) KS domains, respectively (Fig. 1e, f). The second mPKS pair contains six (c) and two (d) KS domains, respectively (Fig. 1e, f). The longest of the mPKS genes in each species is *mPKSa* and *mPKSc*. *Eta-mPKSa* and *c, Ed-mPKSc*, and *Ep-mPKSc*, all encode proteins approximately 10,000 amino acids long (*Ep-mPKSa* is reduced in sized, discussed below). These are the longest predicted proteins in the genomes of each species.

The *Eutiarosporella* mPKS genes are in the middle of a scaffold, surrounded by genes of fungal origin. However, to corroborate the findings qRT-PCR expression analysis was performed on the modular pair from *E. tritici-australis*. Expression of the genes from *E. tritici-australis* were tested under three nutrient substrate conditions: potato dextrose broth (PDB), fries medium (FM), and on wheat. Expression of the two genes was observed under all three conditions, and compared in relation to expression of beta-tubulin (Supp. Fig. 1).

To determine whether any other fungal species possess mPKS genes, we performed a Blastp analysis of the mPKS sequences against the NCBI or JGI databases. This revealed that the only other fungal homologues of these genes were identified in *M. phaseolina*, a close relative, Botryosphaeriaceae spp.. Three mPKS genes were predicted for *M. phaseolina*, however, after re-annotation with FGENESH+ (softberry.com), these were reduced to one pair that share a 5’ non-coding region and are predicted on opposite strands (*Mp-mPKSa* and *Mp-mPKSb*) (Fig. 1d). The encoded amino-acid sequences contain six (a) and two (b) KS domains, respectively (Fig. 1e, f). To the best of our knowledge, mPKS genes have previously never been observed in fungi. It is also significant to note that no other fungi represented on these publically available databases harbour genes encoding mPKSs. Therefore, the Botryosphaeriaceae mPKS genes described in this study are the first observed fungal mPKS genes. As mentioned above, not all of the KS domains encoded by the predicted Botryosphaeriaceae mPKS genes are followed by an acyl transferase (AT) domain (Fig. 1e, f blue text). AT domains, which are the functional domain responsible for substrate loading in PKS systems is considered one of the minimal PKS domains along with KS and acyl carrier protein (ACP) domains and hence usually present in each PKS module (Fig. 1e,f). However, among the *Eutiarosporella* mPKS genes, AT domains only follow *Eta-*KSa2 and 4, *Eta-*KSb1, *Ed-*KSa4, *Ed-*KSb1, *Ed-*KSc2 and 4, *Ep*-KSb1, *Ep-* KSc2 and 4, *Ep-*KSd1, *Mp-*KSa2 and 4, and *Mp-*KSb1. *Ep-mPKS-a* is not predicted to encode any AT domains (Fig. 1e, f). Each of the four species possesses at least one longer mPKS gene (either a or c) and one shorter mPKS gene (either b or d) that share a generally conserved domain structure with the other species (Fig. 1e,f).

### Botryosphaeriaceae mPKS genes were likely obtained via horizontal gene transfer

To investigate the evolutionary origins of the mPKS genes in the WGD *Eutiarosporella spp.*, a maximum likelihood (ML) phylogeny was performed comparing the KS domains from the mPKSs to other predicted PKS proteins (non-modular) within their genomes. Additionally, the KS domains from close BLAST hits to the mPKS*-*KS domains and to KS domains from known fungal and bacterial PKS genes (Fig. 2, Supp. Fig. 2), both iterative and modular, were included. The phylogeny demonstrated that the KS domains from the *Eutiarosporella spp.* and *M. phaseolina* mPKSs resolve separately from other fungal PKSs within a clade containing bacterial and protist PKSs (Fig. 2, Supp. Fig. 2). The KS domains from fungal iPKSs and the KS domains from the Botryosphaeriaceae mPKSs appear to have distinct evolutionary histories, whereby KS domains from this newly discovered group of mPKSs do not group with the fungal KS domains from iterative PKSs. This data indicates a potential bacterial or protist origin (Figure 2).

**Figure 2:**
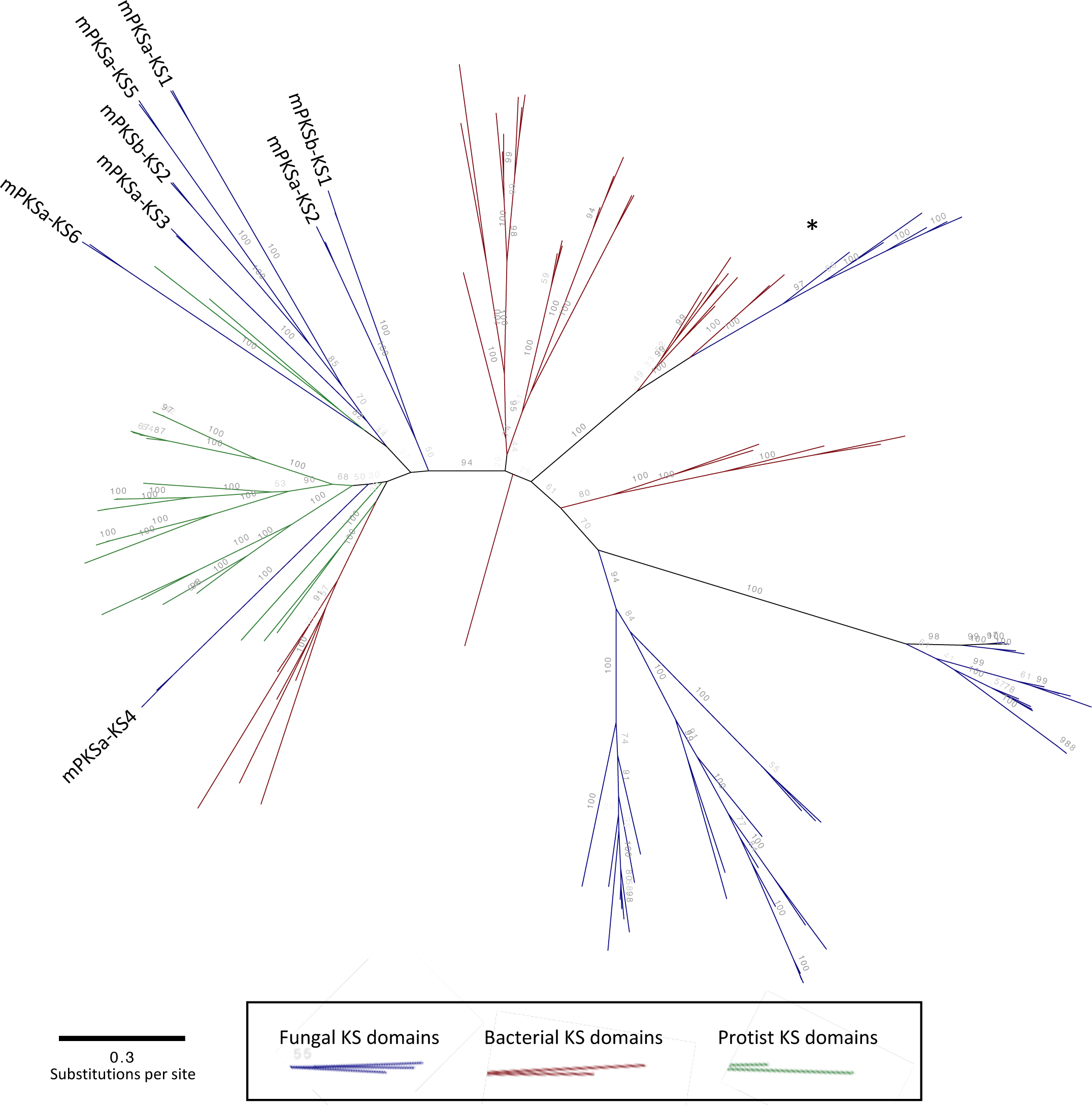
A maximum likelihood phylogeny comparing KS domains from *E. tritici-australis’* and *M. phaseolina’s* mPKSs, other fungal PKSs, protist PKSs, and bacterial PKSs. (*) Highlights 6MAS/mellein synthase KS domains from fungi and bacteria. Bootstrap support values below 70% are faded to reduce noise.

We constructed a second phylogenic tree to assess whether horizontal gene transfer is a likely scenario for the origin of the mPKS genes using the same alignment (Supp. Files). This tree was built using RaxML with the addition of a constraint file, constraining bacterial and protist KS domains together, and the Botryosphaeriaceae mPKS KS domains iterative fungal KSs. KS domains from both bacterial and fungal 6MAS/mellein synthase sequences were not constrained. These genes in fungi have previously been described as horizontally acquired from bacteria (Schmitt and Lumbsch, 2009), and in our initial tree both fungal and bacterial sequences clade together (clade represented by (*): Fig. 2, Supp. Fig. 3). As such they were allowed to move unconstrained during the tree build to prevent these sequences corrupting the statistical analysis, described below. The best constrained tree was compared to the best tree from the initial tree where no clade constraints imposed (Each shown in Supp. Fig. 3), using Consel with the Kishino-Hasegawa (KH) and Shimodaira-Hasegawa (SH) tests (Shimodaira and Hasegawa, 2001). The alternative topologies were determined as statistically different (p<0.05) from the unconstrained trees with scores of p<0.034 for both the KH and SH tests (Supp. Files) with the unconstrained tree being more likely. These results support a hypothesis that the Botryosphaeriaceae mPKSs were obtained via horizontal gene transfer (HGT). Due to lack of clear phylogenetic resolution between the fungal mPKSs and any other organism, it was not possible to determine a potential source species of the Botryosphaeriaceae mPKS genes (Fig. 2, Supp. Fig. 2). In addition, no other non-fungal species share a similar domain structure to the Botryosphaeriaceae mPKSs.

### mPKS c and d in *E. darliae* and *E. pseudodarliae* arose via a gene duplication event

As described earlier, *E. darliae* and *E. pseudodarliae* each possess additional copies of the mPKS genes. In *Supplementary Figure 4* it is clear that the individual KS domains from the mPKSa are more similar to the KS domains in mPKSc at the respective corresponding position than they are to the other KS domain within its own protein (demonstrated for all predicted KS domains in Supp. Fig. 4). Likewise, the individual KS domains from mPKSb and mPKSd are also more closely related to each other than they are to other KS domains within the protein. Each of the clades containing the KS domain groupings is strongly supported (100% bootstrap support). Based on the phylogenetic data we conclude that the additional pair of mPKS genes in *E. darliae* and *E. pseudodarliae* are the result of a duplication event (Supp. Fig. 4).

The only KS domains that did not follow the above described pattern were *Ep-* mPKSa-4 and Ep-mPKSa-5, which resolved with the other mPKSa/c KS domains 5 and 6, respectively (Supp. Fig. 4). *Ep-*mPKSa only harbours five KS domains instead of six KS domains predicted in the other mPKSa and c proteins (Fig. 1e). This indicates that the KS domain in the fourth position for mPKSa/c from *E. darliae, E. tritici-australis*, and *M. phaseolina*, and in *Ep-*mPKSc, has been lost from *Ep-*mPKSa. Inclusive of missing the fourth position KS domain, there is a significant size difference between *Ep-mPKSa* and *Ep-mPKSc*. The entire predicted gene length of *Ep-mPKSa* is only 17322bp and the predicted amino acid sequence is only 4342aa, whereas, *Ep-mPKSc* is 33137bp, and the predicted amino acid sequence is 9935 aa. Likewise, *Ed-mPKSa* is shorter than *Ed-mPKSc*, however, the difference is far less. *Ed-mPKSa* is 29933bp, with a predicted amino acid length of 9264aa, in contrast to *Ed-mPKSc*, which is 31814bp in length (10090aa). In *E. darliae*, we did not identify a second copy of the shorter mPKS (*mPKSb* and *d*). We predict that if *mPKSa*/*c* and *mPKSb/d* were duplicated together, it was likely that a hypothetical *Ed-mPKSd* was subsequently lost. Combined, this data indicates that after the duplication event, a number of deletion mutations resulted in loss of functional PKS domains.

We compared gene synteny surrounding *E. darliae* and *E. pseudodarliae’s* mPKS gene regions with the genes surrounding *E. tritici-australis*. The two genes downstream of *Eta-mPKSa* were conserved downstream of all *Eutiarosporella mPKSa* and *mPKSc* genes (Fig. 1). The encoded protein sequence of the gene immediately adjacent to mPKSa has a predicted serine hydrolase domain (pfam03959). The encoded protein sequence for the second gene had close BLASTp hits among the Botryosphaeriaceae, but no predicted domain. Of the two genes, *M. phaseolina* has an unannotated DNA sequence, in opposite orientation with sequence similarity to the putative serine hydrolase. (Fig. 1d) However, there is no evidence for the second gene on this *Mp*-mPKS scaffold.

The genes downstream of each of the *mPKSc* and *d* genes were not conserved across all species (Supp. Fig. 5) However, a syntenic relationship was established between *Eta-*m*PKSa* and *b* and, *Ep* and *Ed-mPKSa* and *b* (Supp. Fig. 5). Based on analysis of re-arranged, syntenic gene-blocks, we hypothesize that *Ep-mPKSa* and *b*, and *Ed-mPKSa* and *b* were the original *mPKS* pair in each of their respective genomes (Supp. Fig. 5). For this prediction to be accurate, the duplication of *mPKSa* and b to form *mPKSc* and *d* must have occurred after speciation of *E. tritici-australis* from the more closely related *E. darliae* and *E. pseudodarliae*, i.e. *E. tritici-australis* never possessed additional mPKS pair c and d.

### *E. darliae* and *E. pseudodarliae* possess a SM cluster with multiple hybrid PKS-NRPS genes, which was lost from *E. tritici-australis*

We searched additional examples of SM cluster expansion within the WGD *Eutiarosporella spp.*, similar to the duplicated mPKS genes within *E. darliae* and *E. pseudodarliae*. Examining the SM clusters identified via antiSMASH, we found that both *E. darliae* and *E. pseudodarliae* possess a SM cluster, within which there are two PKS-NRPS genes (*Hybrid-1* and *Hybrid-2*) (Fig. 3a). In contrast, *E. tritici-australis* is not predicted to encode any PKS-NRPS genes (Fig. 3a, Table 2). Both *Hybrid-1* and *Hybrid-2* are predicted to encode amino acid sequences of similar length and domain structure (Fig. 3b). As above, we performed a phylogenetic analysis comparing the KS domains of these against other close BLAST-hits from NCBI. *Hybrid-1* and *Hybrid-2* resolved in separate clades, indicating that they are not duplicate genes (Fig. 3b).

**Figure 3:**
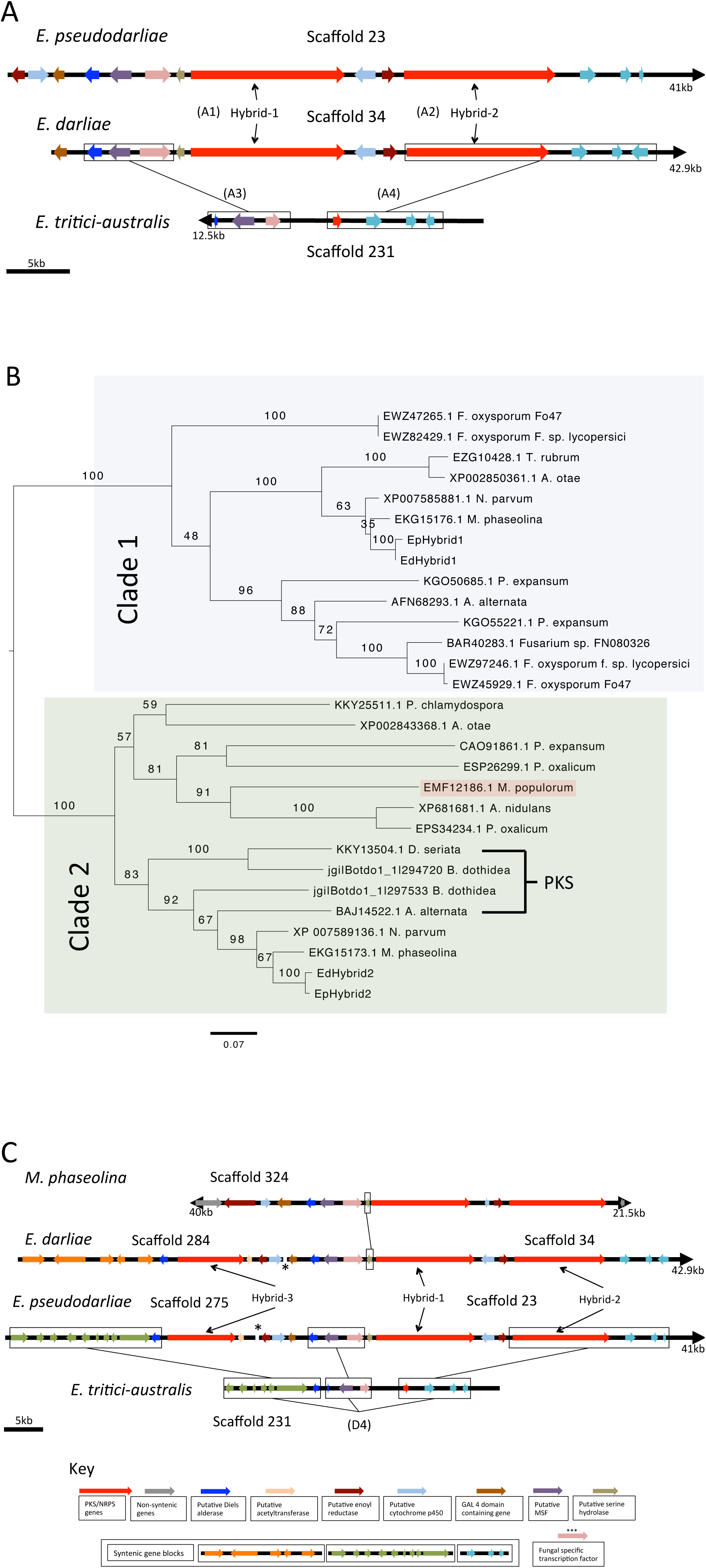
*E. darliae* and *E. pseudodarliae* each harbour a SM cluster with multiple hybrid *PKS/NRPS* genes (*Hybrid-1, 2 and 3*), which was lost in *E. tritici-australis*. A) A graphical representation of *Hybrid-1* and *Hybrid-2* from each of *E. darliae* and *E. pseudodarliae* on a single scaffold, compared with the syntenic gene region in *E. tritici-australis*. B) A maximum likelihood phylogeny comparing KS domains of *Hybrid-1* and *2*, as well as close BLAST-hits from the NCBI non-redundant database. *Hybrid-1* and *Hybrid-2* resolved separately on this phylogeny ((Clade 1) and (Clade 2), respectively). A *PKS-NRPS* of note, harboured by a poplar pathogen, *M. populorum* (gi|453084141), and which resolves in *Hybrid-2’s* clade, is highlighted in red. Also noted are four KS sequences belonging to putative PKSs, not PKS-NRPSs. C) Graphical representation of the entire predicted cluster of *Hybrid-1, 2*, and *3*, in *E. darliae* and *E. pseudodarliae* across two scaffolds, and the syntenic genes present in *E. tritici-australis* that are on a single scaffold. A representation of *M. phaseolina’s Hybrid-1* and *2* genes are included to demonstrate that they are also co-localised, however, not *M. phaseolina’s Hybrid-3* homologue. Outlined serine hydrolase between *M. phaseolina* and *E. darlie* highlights difference between the two clusters. Highlights difference between the *<u>Eutiarosporella spp.</u>* <u>and *M. phaseolina’s* cluster</u>. (*)Highlights scaffold overhang joint by two syntenic genes linking the cluster’s scaffolds in *E. darliae* and *E. pseudodarliae*. Key: describes genes present in each cluster, as well as syntenic gene-blocks.

Several known secondary metabolite related-genes are found neighbouring *Hybrid-1* and *Hybrid-2*. *E. pseudodarliae* has nine genes: seven of these genes are upstream of *Hybrid-1*: an enoyl reductase-like gene (cd08249), a cytochrome p450 (pfam00067) a GAL4 domain containing gene (smart00066), a Diels-Alderase-like gene, an MFS protein (cd06174), a fungal specific transcription factor (FSTF), and a serine hydrolase (pfam03959) (Figure 3c). Two more genes are upstream of *Hybrid-2*: an enoyl reductase-like gene and a cytochrome p450 gene. *E. darliae* has seven of the nine genes listed above, in the same order but is missing the first two genes due to the end of the scaffold. We identified these two genes missing from *E. darliae’s* cluster on a separate scaffold (Figure 3c, Scaffold 284). In addition to these two genes, this scaffold contained a third NRPS gene flanked by a putative acetyltransferase gene and a putatative Diels alderase gene. For consistency among genes in this cluster, we define this NRPS as *Hybrid-3*. *E. pseudodarliae* also possesses these other three genes, on a separate scaffold (Figure 3c, Scaffold 275). In both *E. darliae* and *E. pseudodarlieae* the ends of these scaffolds form short overlaps indicating that these three synthase genes (*Hybrid 1-3*) form one large SM cluster (Fig. 3c).

We compared the genes from these clusters to their syntenic positions in the two species with best-BLAST homologues, *N. parvum* and *M. phaseolina* (Table 2). *M. phaseolina* possesses a cluster that contains homologues of *Hybrid-1* and *Hybrid-2*, as well as the initial nine associated genes (Fig 3c). *N. parvum* also possesses homologues of these proteins, however, they are not located within a single cluster (Supp. Fig. 6). *Hybrid-3* on the second *Eutiarosporella* scaffold has a best-BLAST-hit on NCBI with a predicted PKS-NRPS from *N. parvum* (*NRPS-*4; Table 2, Supp. Fig. 7a). A homologous protein is also found in *M. phaseolina*, not co-located with the other two PKS-NRPS genes (Supp. Fig. 7a), In the *Eutiarosporella spp.*, the N-terminal region *of Hybrid-3* has been truncated, removing what are the KS domains found in *N. parvum* (Supp. Fig. 7b).

We could not identify intact copies of *Hybrid-1, -2*, or *-3* in *E. tritici-australis*. tBLASTx searches identified a predicted partial blast-hit to *Hybrid-2* (681bp, e-value 7.12e-86, 72.7% pairwise identity) (Fig. 3a, c). Genes found flanking *Hybrid-1*, *-2*, and *-3* in *E. darliae* and *E. pseudodarliae* were used as tBLASTx queries to search *E. tritici-australis*. This revealed the presence of some of these genes in *E. tritici-australis*, on the same scaffold as the partial *Hybrid-2* (Fig. 3a,c). Secondary metabolite related genes identified include: *Hybrid-1’s* Diels-Alderase-like, MSF, and serine hydrolase genes, and *Hybrid-3’s* upstream Diels-Alderase-like gene. Additional syntenic genes identified, that were not predicted to encode predicted SM associated domains, include: three homologues to genes downstream of *Hybrid-2* in *E. darliae* and *E. pseudodarliae* and six homologues to genes upstream of *E. pseudodarliae’s Hybrid-3*. The presence of all of these genes on a single scaffold in *E. tritici-australis*, provides further support that *Hybrid-1, -2*, and *-3* are a part of a larger contiguous cluster (Fig. 3c).

### *E. darliae* and *E. pseudodarliae* are able to induce disease symtoms in a woody-plant species

A number of PKS-NRPS sequences that resolve within the clade containing *Hybrid-2* (Fig. 3b) are harboured by pathogens of woody-plants. For example, *Botryosphaeria dothidea* (jgi|Botdo1_1|294720 and jgi|Botdo1_1|297533), *Neofusicoccum parvum* (gi|615428986), *Diplodia seriata* (gi|821057639), *Alternaria alternata* pathovar tangerines (gi|302562829), *Mycosphaerella populorum* (gi|453084141), and *Phaeomoniella chlamydospora* (gi|821070483). In particular, two of these proteins are linked to facilitation of pathogenicity in woody plants (Miyamoto *et al*., 2010, Dhillon *et al*., 2015). The first is the protein from *Alternaria alternata* pathovar tangerines (gi|302562829, BAJ14522.1; BLASTp with *E. pseudodarliae’s Hybrid-2’s* KS domain: e-value 0.0, 81% identity). This protein, called ACTTS3, is a PKS used in the production of ACT-toxin, a host-specific toxin that enables infection of tangerines (Kohmoto *et al*., 1993). Despite resolving among PKS-NRPS genes (Fig. 3b), ACTTS3 lacks the NRPS domains. ACTTS3 resolves in a monophyletic clade with Botryosphaeriaceae PKS-NRPS genes, three of which also lack NRPS domains (*B. dothidea*: JGI|Botdo1_1|294720 and JGI|Botdo1_1|297533, and *D. seriata* gi|302562829) (Fig. 3b).

The second example is in the poplar-pathogen, *Mycosphaerella populorum* (gi|453084141, EMF12186.1; BLASTp with *E. pseudodarliae’s Hybrid-2’s* KS domain: e-value 0.0; 67% identity). Horizontal acquisition of this PKS-NRPS gene cluster is predicted to be a factor differentiating disease severity between itself and a close relative, *M. populicola*, that lacks the cluster (Dhillon *et al*., 2015). The PKS-NRPS and clustered genes are up-regulated on wood-substrate (Dhillon *et al*. 2015). *M. populorum’s* PKS-NRPS gene cluster only shares a limited syntenic relationship with *E. darliae* and *E. pseudodarliae’s Hybrid-2’s* cluster*s* (Supp. Fig. 8). Conserved with *M*. populorum’s cluster are both *Hybrid-2’s* enoyl reductase-like gene (comparative tBLASTx e-value 1.90e-99) and cytochrome p450 gene (comparative tBLASTx evalue 1.90e-99) (Supp. Fig. 8). In addition, using the amino acid sequence encoded by *M. populorum’s FSTF* gene as a tBLASTn query, we identified a homologous stretch of DNA, on a different scaffold to *Hybrid-2* (e-value 1.07e-38). In comparison, *Hybrid-1’s FSTF* gene was a less significant BLAST-hit (e-value 4.90e-07). No gene was predicted at the best-BLAST-hit site in *E. pseudodarliae*, and it is only a partial hit (1065bp vs. 2053bp for *M. populorum’s* complete *FSTF*) (Supp. Fig. 8).

We hypothesized that the WGD *Eutiarosporella spp*. that possess homologues of the *Hybrid* genes might induce disease in woody plants. To test this hypothesis, attached leaf assays were performed on various woody plants, using all three *Eutiarosporella spp.*. Leaves were cut prior to inoculation to facilitate fungal penetration. Disease symptoms attributed to *E. darliae* and *E. pseudodarliae* were particularly pronounced in *Hakea salicifolia*, and so we use this plant species to represent the ability of the WGD *Eutiarosporella spp.* to induce disease (Fig. 4, Supp. Figs. 9 and 10). After three days post-inoculation (DPI) necrotic lesions and disease symptoms were observed in leaves inoculated with *E. darliae* and *E. pseudodarliae* (Fig. 4, Supp. Figs. 9 and 10). In contrast, almost no disease symptoms were evident on leaves inoculated with *E. tritici-australis* (Fig. 4, Supp. Figs. 9 and 10). Attached leaf disease assays were re-performed using an additional isolate of each of *E. darliae* and *E. pseudodarliae* as well five additional isolates of *E. tritici-australis* (Supp Fig. 11). Extensive necrotic lesions occurred 3DPI in leaves inoculated with the former two species, however, little to no necrosis was induced beyond the site of inoculation by any of the isolates of *E. tritici-australis* (Supp. Fig. 11). *E. darliae* and *E. pseudodarliae* were re-isolated from diseased tissue, away from the inoculation site, thereby satisfying Koch’s postulate. Expression of *Hybrid-1*, *Hybrid-2*, *Hybrid-3* and the cluster’s putative *TF* gene are up-regulated in *E. darliae* when the fungus is grown on Hakea wood compared to when growth in potato dextrose broth (PDB) (Supp. Fig. 12). Details of disease progression and comparisons among each of the three species are displayed in more detail in the supplementary data (Supp. Figs. 9 and 10).

**Figure 4:**
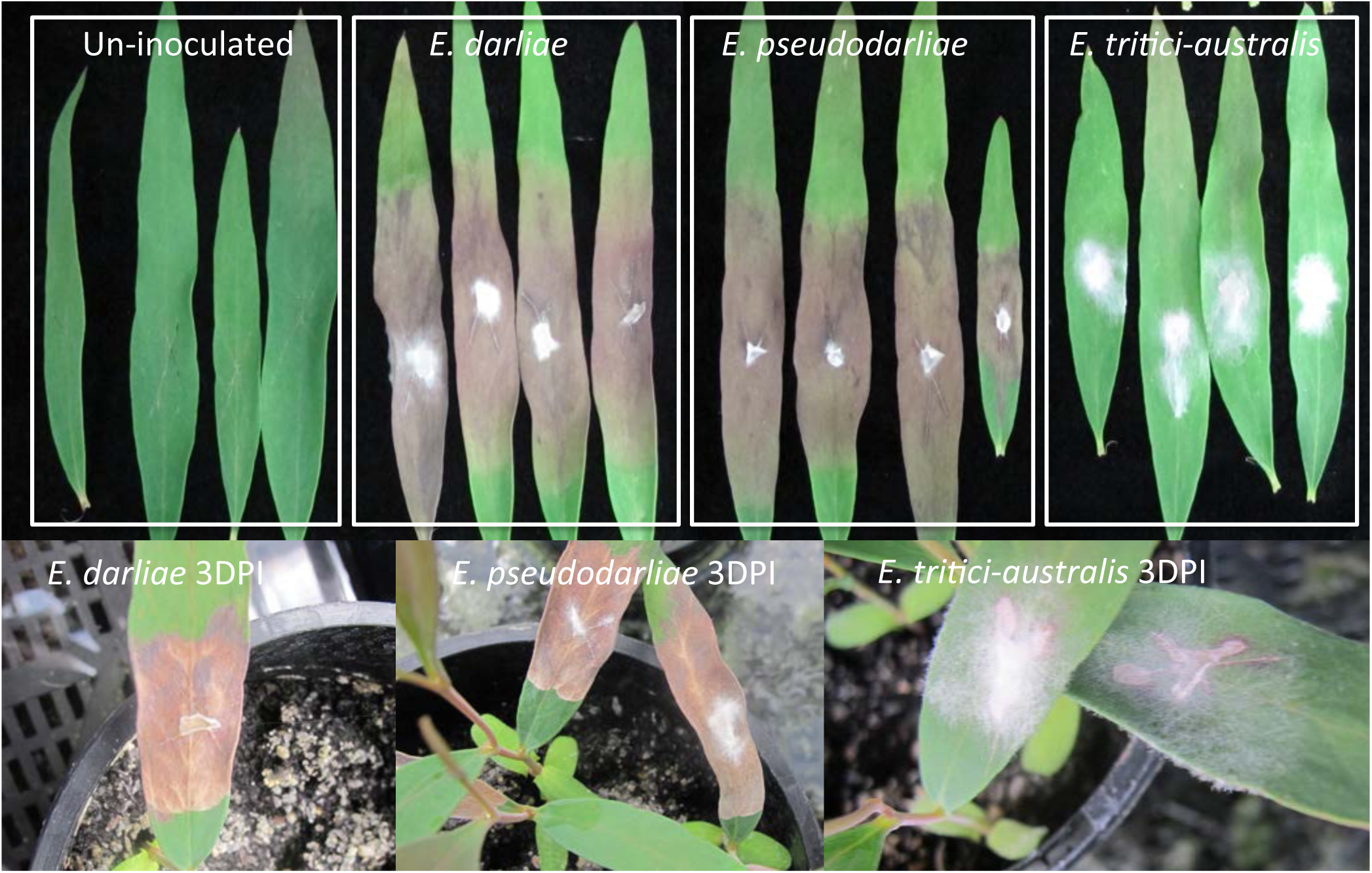
*E. darliae* and *E. pseudodarliae* induce necrotic disease symptoms in *Hakea salicifolia*, whereas *E. tritici-australis* does not. Detached leaf (upper panels) and attached leaf (lower panels) assays of necrosis development on *H. salicifolia*. Wounded leaves of were inoculated with the WGD *Eutiarosporella sp..* The un-inoculated leaves displayed no disease symptoms. Leaves inoculated with both *E. darliae* and *E. pseudodarliae* displayed significant necrotic symptoms spreading from the cut site at three days post inoculation (3DPI). Leaves inoculated with *E. tritici-australis* do not display significant disease symptoms beyond the site of inoculation at 3DPI. Note: the detached leaf assay panels with *E. pseudodarliae* and *E. tritici-australis* were beneath the negative control and *E. darliae* panels in the original image, but were cut-and-pasted to be side-by-side.

## Discussion

Gene acquisition and loss are important adaptation events in the evolution of fungal genomes. We highlight the occurrence of these events relating to secondary metabolite (SM) genes in the genomes of each of the WGD *Eutiarosporella spp*. First, we described the acquisition of modular PKSs (mPKSs), the first of their kind reported in fungi. Second, we present that an SM cluster comprising of three hybrid PKS/NRPS genes present in *E. darliae* and *E. pseudodarliae* were lost from *E. tritici-australis*, and that this loss likely impacted the latter species’ ability to infect woody plant species. Combined, these observations illustrate that, despite these fungi having the smallest genomes among the Dothideomycetes, assembled to date, significant genomic adaptation events have still shaped their evolution.

*iPKSs* are ubiquitous throughout fungi and bacteria (Throckmorton *et al*., 2015, Yun *et al*., 2015, Chooi and Tang, 2012, Keller *et al*., 2005), whilst, mPKSs have previously only been described in bacteria and protists (Khosla, 1997, John *et al*., 2008, Zhu *et al*., 2002). In bacteria, examples of SMs produced by mPKSs include antibiotics, such as erythromycin and spiramycin (Khosla, 1997). mPKSs were only recently observed in protist species (John *et al*., 2008, Zhu *et al*., 2002). As of yet, a definitive link between protist SMs and PKSs has not been made due to the difficulty involved in culturing and genetically manipulating these organisms (Kohli *et al*., 2015). The presence of mPKSs within the WGD *Eutiarosporella*, previously undescribed in fungi, is an exciting advance in our knowledge of the diversity of fungal secondary metabolism. This finding is consistent with others that report “bacterial-only” SMs in newly sequenced fungal species. For example, a limited number of instances of Type-3 PKSs in fungi were observed, which were thought to be limited to plants and bacteria (Hashimoto *et al*., 2014). Similarly, a recent study found that in addition to using hybrid PKS (N-terminal)-NRPS (C-terminal) proteins, various fungi also use hybrid NRPS (N-terminal)-PKS (C-terminal) proteins for SM biosynthesis; these too were previously thought to be limited to bacteria (Yun *et al*, 2015). Our discovery of fungal mPKS genes further demonstrates that the boundaries between fungal and bacterial SM biosynthesis are not clear-cut.

Additional unanswered questions remain, for example, what is the mechanism these proteins use to synthesize SMs? It is generally considered that the minimal module required for a Type-1 PKS consists of a KS domain (extension unit) and an acyl transferase (AT) domain (substrate loading domain) and acyl carrier protein (ACP) (Chooi and Tang, 2012, Hertweck, 2009, Keller *et al*., 2005, Throckmorton *et al*., 2015). However, not all Botryosphaeriaceae mPKS genes’ modules conform to this structure, as some lack AT domains. Lack of these domains does not necessarily impact the ability of the synthase to function. There are functional AT-less PKS proteins with modules that are complemented by separately encoded AT domains (Cheng *et* al., 2003, Lohman *et al*., 2015) and there are also SM synthases that produce a main product, and then additional, different metabolites by skipping over particular modules (Wenzel *et al*., 2005). Elucidation of produced metabolites will begin to shed light on how these proteins function.

In addition to elucidating the mPKS products, we similarly intend to determine the product(s) of the three co-localised *PKS-NRPSs* (*Hybrid-1, 2*, and *3*). As *Hybrid-1*, *-2*, and *-3* were lost from *E. tritici-australis*, we assume they are not major virulence factors for these fungi on wheat. Instead, our interest stemmed from the evolutionary implications associated with harbouring these genes. This led us to examine these species’ ability to infect woody plants in a reverse ecological manner (Ellison *et al*., 2011). Based on our observations, we hypothesise that woody-plants might act as an alternative host-reservoir for the fungi. Perhaps inoculum to infect wheat originates from fungal growth in woody plants. Very little is known about the lifestyle of these fungi beyond infection of wheat. The majority of Botryosphaeriaceae spp. are reported as pathogens or endophytes (and opportunistic pathogens) of woody plants (Slippers and Wingfield, 2007). Similarly, other *Eutiarosporella spp.* have been isolated from woody plants (Jami *et al*, 2012, Jami *et al*., 2013). To develop a better understanding of the lifestyle of these pathogens it would be beneficial to increase sampling of non-agricultural hosts (both monocots and dicots) for fungi growing undetected (Hyde *et al*., 2010).

Gene loss and rapid acquisition of new genes (i.e. evolution of new orphan genes or HGT) are two sides of the same coin in a pathogen’s ability to adapt to new environments and hosts. In this regard, *E. tritici-australis’* loss of the hybrid PKS-NRPS cluster is akin to the example of *M. populorum’s* horizontal acquisition of a homologous cluster from *Penicillium oxalicum* (Dhillon *et al*., 2015). What drove *E. tritici-australis* to lose its PKS-NRPS cluster remains unknown. Our preliminary data suggests the presence of these genes could be advantageous for a pathogenic lifestyle in woody plants, and so if this species has made a committed jump to life in wheat, their loss may not be a disadvantage. Another possibility for why *E. tritici-australis* lost this cluster is that it became unfavourable to keep these genes. A similar scenario was depicted for an *M. oryzae PKS-NRPS* cluster, *ACE1* (Khaldi *et al*., 2008, Berruyer *et al*., 2003). The SM synthesised by *ACE1’s* encoded proteins is recognized by rice cultivars harbouring the receptor gene, *Pi33*. Recognition of the *ACE1* product by *Pi33* leads to a strong defence response from the plant, there-by putting selective pressure on the pathogen to lose or stop making the SM (Berruyer *et al*., 2003). If the product of the WGD *Eutiarosporella spp.’* cluster is recognized by a particular host, then perhaps this is why it has been lost in *E. tritici-australis*. It is pertinent to note that in South Australia, *E. tritici-australis* has been observed as the most virulent of the three species on wheat (personal communications with Marg Evans, SARDI). Further work studying host-range and host responses to all three WGD species would have to be performed to determine any potential link between these hybrid PKS-NRPS and putative resistance in grasses.

## Conclusion

The discovery of horizontally acquired mPKS genes in the WGD *Eutiarosporella spp*. and *M. phaseolina* expands the known boundaries of fungal SM biosynthesis capabilities, as these are the first described examples of mPKS genes in fungi. This is a significant finding for the field of fungal SM biosynthesis and also provides a fascinating target for future research endeavours. The observation of a PKS-NRPS gene cluster harboured in *E. darliae* and *E. pseudodarliae*, but lost in *E. tritici-australis*, was significant to our research through increasing our understanding of these three pathogens’ lifestyles. This is because it led us to explore the variation in host-range and virulence among these fungi, which demonstrated that *E. darliae* and *E. pseudodarliae*, but not *E. tritici-australis*, are able to induce disease in examined woody plants. Further, we speculate that perhaps these fungi use woody plants as secondary hosts, which would have implications on potential efforts to manage the disease. Combined, these observations relating to the secondary metabolite capabilities of the WGD *Eutiarosporella spp*. demonstrate that rapid genomic adaptations have played an important role in the evolution of these three, fungal species.

## Materials and methods

### WGD fungi sequencing, assembly and annotation

Genome assemblies are described in Thynne *et al*., (2015b) for each of *Eutiarosporella tritici-australis, Eutiarosporella darliae*, and *Eutiarosporella pseudodarliae* (Thynne *et al*., 2015). Illumina paired-end sequencing was performed using a HiSeq™ 2000 (Illumina, USA) at the John Curtain School of Medical Research, Australian National University (ANU). The resulting sequenced reads were quality trimmed using Trimmomatic v0.27 (Lohse *et al*., 2012) and assembled *de novo* with SPAdes v2.5.0, with –k mer values 21, 33, 55, and 77 (Bankevich *et al*., 2012). Gene sets for each species were predicted and annotated with the MAKER genome annotation pipeline with the gene sets from *Botryosphaeria dothidea* and *Parastagonospora nodorum* used as training genes. (Cantarel *et al*., 2008). Where required, genes were re-annotated with FGNESH+ on the online server at softberry.com (**Solovyev, 2001)**. Annotations were visualized within Geneious v7.1.8 (Biomatters, New Zealand). Secondary metabolite genes were predicted with the online antiSMASH (v.2) server (Weber *et al*., 2015b). Protein domains were predicted with Interproscan v.1.0.6. (Quevillon *et al*., 2005).

### Genome analysis of *Eutiarosporella* and *Macrophomina* modular PKSs and the hybrid PKS-NRPS cluster

The relative genome-positions of the modular PKS genes (mPKS), hybrid *PKS-NRPSs*, and surrounding genes were determined in Geneious v7.1.8, through manual analysis of the containing assembled scaffolds. Annotated genes were used as BLASTn and tBLASTx queries to search for these genes and compare synteny on the genome-scaffolds of the alternative *Eutiarosporella spp.*.

### Phylogenetic analysis and alternative topology testing of *Eutiarosporella* KS domains

The ketoacyl synthase (KS) domains for the predicted polyketide synthase (PKS) amino-acid sequences from each of the wheat-infecting *Eutiarosporella spp.* were used as BLASTp (Altschul *et al*., 1990) queries to search the NCBI non-redundant (NR) database for close homologues. Similarly, the KS domain of *Penicillium aethiopicum* VrtA amino acid sequence and the *Alternaria alternata* PKSj amino acid sequence were used to search for additional fungal PKS homologues. Selections of these PKS homologues were downloaded from NCBI and the protein domains for each predicted with Interproscan v.1.0.6. (Quevillon *et al*., 2005). The predicted KS domains from all of the sequences were extracted and aligned with MUSCLE v3.5 (Edgar, 2004) and the internal Geneious aligner (Geneious v7.1.8 (Biomatters, New Zealand)) (Supplementary file 1). *Ep-*mPKSa KS1,2, and 3, and *Ed*-mPKSa KS1 and 2, were removed from this tree due to decreased amino acid sequence length (these were included in a secondary tree comparing the *Botryosphaeriaceae* mPKS-KS domains). A maximum likelihood tree with 500 bootstraps was created using RaxML v8 (Stamatakis, 2014) with the rapid bootstrapping command “-f a”, the random number seed and starting tree commands of “-p 1234” and “-x 1234”, respectively.

To test the alternative topology, another tree was generated using a new alignment without mellein/6-MAS sequences (Supplementary file 2). A further tree was generated with the addition of the RaxML multi-furcating tree command “-g” with a kingdom segregated constraint file. The “best-tree” output from both the constrained and unconstrained trees were concatenated and used as the input file to generate a site-log likelihood file from RaxML using the “-f g” command. The output file in treepuzzle format was fed into Consel (Shimodaira and Hasegawa, 2001) to perform Kishino-Hasegawa (KH) and Shimodaira-Hasegawa (SH) tests to compare the significance of the two trees’ topologies.

### Phylogenetic analyses of KS domains from the hybrid *PKS-NRPS* genes

The KS domains from the encoded amino acid sequences of the hybrid *PKS-NRPS* genes were used as BLASTp queries to search the NCBI non-redundant (NR) database and the JGI MycoCosm portal for close homologues. Some of predicted protein sequences were shorter than expected (EKG15258.1, KKY13504.1, and were re-annotated using using FGNESH+ on the online server at softberry.com. The domains for each homologue were predicted with Interproscan v.1.0.6. (Quevillon *et al*., 2005) and the online antiSMASH (v.2) server (Weber *et al*., 2015b). Images used were generated in Geneious v7.1.8 (Biomatters, New Zealand) and the online antiSMASH (v.2) server (Weber et al., 2015b). A maximum likelihood tree with 500 bootstraps was created using RaxML 7.2.8 (Stamatakis, 2014) in Geneious v7.1.8 (Biomatters, New Zealand) with the “rapid bootstrapping and search for best-scoring ML tree” command selected. The tree was visualised and annotated in FigTree v1.4.0 (http://tree.bio.ed.ac.uk/software/figtree/).

### Infection assays

Leaves of a variety of woody plants were obtained from plants from the Australian National University campus and surrounding suburbs. *Hakea salicifolia* was used for infection assays in the study. Leaves were wounded in an “X” pattern (approximately 1-2cm) with a scalpel. Wound sites were inoculated with agar plugs covered in growing mycelia and incubated in high-humidity. Disease symptom development was observed over the following week.

Both seedlings and leaves from mature plants were used. To grow the seedlings, *H. salicifolia* seeds were obtained from seedpods from a mature plant. To access the seed, pods were heated at 60°Celsius for six hours (or until open). Seeds were germinated in seed raising mix (The Scotts Miracle-Gro Company, USA). When established shoots appeared, the seedlings were further fertilized with Osmocote (The Scotts Miracle-Gro Company, USA).

## Acknowledgements

ET was supported by a Grains Research and Development Corporation Graduate Research Scholarship. YHC is an Australian Research Council (ARC) Future Fellow. The authors also acknowledge the Australian Plant Phenomics Facility which is supported under the National Collaborative Research Infrastructure Strategy of the Australian Government

---

**Supplementary Figure 1: Relative expression of the *Eta-mPKSa* and *Eta-mPKSb* grown on wheat, fries medium, or potato dextrose agar.** Error bars represent standard error.

**Supplementary Figure 2: Expanded layout of a maximum likelihood phylogeny comparing KS domains from *E. tritici-*australis and *M. phaseolinas’* mPKSs against KS domains from fungi, bacteria, and protists.** Botryosphaeriaceae mPKS-KS domains are highlighted in dark blue. Bacterial KS domains are highlighted in red. Protist KS domains are highlighted in green. Fungal, iterative PKS-KS domains are highlighted in teal.

**Supplementary Figure 3: Best trees from both the unconstrained phylogenetic analysis and the constrained phylogenetic analysis.**

**Supplementary Figure 4: Maximum likelihood phylogeny comparing all KS domains from all Botryosphaeriaceae mPKSs.** Both (*) denote that *Ed-*mPKSa domains do not follow the usual pattern, demonstrating the loss of the fourth position KS domain.

**Supplementary Figure 5: Gene-block synteny confirms the original of the mPKS pairs in *E. darliae* and *E. pseudodarliae*. *M. phaseolina’s* mPKS pair located on a non-syntenic locus.** A) The gene blocks downstream of the *Ep-mPKSb, Ep-mPKSd*, and *Ed-mPKSb* were non-syntenic to the genes downstream of *Eta-mPKSb*. The four genes downstream of *Ep-mPKSb* and six genes downstream of *Ed-mPKSd* are syntenic with a gene-block in *E. tritici-australis* on distal contig (Block X). The four genes downstream of *Eta-mPKSb* form a syntenic gene-block with four genes on another contig in *E. darliae* and *E. pseudodarliae* (Block Y). Adjacent to Block X in *E. tritici-australis* and adjacent to Block Y in *E. darliae* and *E. pseudarliae* is a syntenic seven gene-block (Block Z). The relationship between the three gene blocks, X, Y, and Z, indicates a gene-rearrangement event swapped these from their original configuration. B) There is a gene-block downstream of *Mp-mPKSa* (Block i) and a gene-block downstream of *Mp-mPKSb* (Block ii). *E. tritici-australis* shares these syntenic gene blocks, which are adjacent to each other (Block i & ii). A link between *E. tritici-australis’* scaffold harbouring Block i & ii and its scaffold harbouring the *mPKS* genes was not determined.

**Supplementary Figure 6: Graphical representation of gene-synteny between *E. pseudodarliae’s Hybrid-1* and *2* harbouring scaffold and the *N. parvum* Scaffolds that each harbour a homologous cluster to either *Hybrid-1* or *Hybrid-2*.** Genes were compared using BLASTn. *Hybrid-1* and upstream genes are syntenic with a *PKS-NRPS* gene cluster on *N. parvum’s* Scaffold 786. *Hybrid-2* and its two adjacent, upstream genes are syntenic with genes on *N. parvum’s* Scaffold 1198. Another, *PKS-NRPS* gene is also present on this Scaffold, removed from the *Hybrid-2-*like gene.

**Supplementary Figure 7: A truncated *PKS-NRPS* was identified in *E. darliae* and *E. pseudodarliae*, related to *Hybrid-1*.** A) Graphical representation of gene synteny across *A. otae, N. parvum*, *E. pseudodarliae*, and *M. phaseolina. E. darliae, N. parvum*, and *M. phaseolina* share syntenic relationship between their *PKS-NRPS* gene and a Diels-Alderase-like gene. B) A nucleotide alignment between *M. phaseolina* and *E. pseudodarliae*. The nucleotide region includes the *PKS/NRPS* genes and the clustered Diels-Alderase-like gene. *E. pseudodarliae* is missing DNA that encodes the N-terminal domains of the *PKS-NRPS* (PKS domains).

**Supplementary Figure 8: A graphical representation of gene synteny between a horizontally acquired *PKS-NRPS* gene cluster in the poplar pathogen, *M. populorum*, and the *Hybrid-2* cluster in *E. pseudodarliae*.** Blue annotations represent genes. Red annotations represent BLAST-hits that are in a region not predicted to encode a gene.

**Supplementary Figure 9: Leaves of growing, young Hakea plants inoculated with mycelia from the WGD *Eutiarosporella spp.*** A) Infection assays three days post inoculation (3DPI) Cut, un-inoculated leaves displayed no necrotic symptoms. Leaves cut and inoculated with *E. tritici-australis* display negligible necrotic symptoms. Leaves cut and inoculated with either *E. darliae* or *E. pseudodarliae* display large necrotic lesions, spreading from the site of inoculation. B) A leaf inoculated with *E. tritici-australis* showing no significant disease progression from 3DPI to 14DPI. C) A leaf inoculated with *E. tritici-australis* at 7DPI with thick aerial mycelia reaching out from the site of inoculation on to the plant’s plastic pot. There appear to be minor signs of chlorosis within the leaf at the site of inoculation. At 14DPI there is only minor increases in necrotic tissue of the leaf, and the aerial mycelia reaching from the plant is wispier.

Supplementary Figure 10: Qualitative and quantitative approximation of necrotic lesion progression in Hakea leaves inoculated with *E. darliae* and *E. pseudodarliae*. A) At three days, post-inoculation (3DPI) there are large necrotic lesions. These lesions spread, changing from a dried brown colour to a brown and black colour with an overlayed powdery, grey hue. B) A graph displaying the lesion spread, in centimetres, away from the site of inoculation, at 3DPI, 5DPI, and 7DPI.

Supplementary Figure 11: Attached leaf disease assays performed on with additional isolates of *E. darliae, E. pseudodarliae*, and *E. tritici-australis*. Photos taken 3DPI. *E. darliae* isolate 2E2 re-used in this assay as a positive control for necrotic lesion development.

Supplementary Figure 12: Relative expression of the hybrid cluster’s *transcription factor, Hybrid-1, Hybrid-2, and Hybrid-3* in *E. darliae* are up-regulated when the fungus is grown on hakea wood as opposed to in potato dextrose broth (PDB). Error bars represent standard error.

## References

Agrios GN. 2005. Plant Pathology. San Diego: Elsevier Academic Press.

Altschul SF, Gish W, Miller W, Myers EW, Lipman DJ. 1990. Basic local alignment search tool. Journal of Molecular Biology 215: 403–410.

Baker SE, Kroken S, Inderbitzen P, Asvarak T, Li BY, Shi L, Yoder OC, Turgeon BG. 2006. Two polyketide synthase-encoding genes are required for biosynthesis of the polyketide virulence factor, t-toxin, by *Cochliobolus heterostrophus*. Molecular Plant-Microbe Interactions 19: 139–149.

Bankevich A, Nurk S, Antipov D, Gurevich AA, Dvorkin M, Kulikov AS, Lesin VM, Nikolenko SI, Pham S, Prjibelski AD, et al. 2012. SPAdes: A new genome assembly algorithm and its applications to single-cell sequencing. Journal of Computational Biology 19: 455–477.

Berruyer R, Adreit H, Milazzo J, Gaillard S, Berger A, Dioh W, Lebrun MH, Tharreau D. 2003. Identification and fine mapping of *Pi33*, the rice resistance gene corresponding to the *Magnaporthe grisea* avirulence gene *ACE1*. Theoretical and Applied Genetics 107: 1139–1147.

Blanco-Ulate B, Rolshausen P, Cantu D. 2013. Draft genome sequence of *Neofussicoccum parvum* isolate UCR-NP2, a fungal vascular pathogen associated with grapevine cankers. Genome Announcements 1: e00339–13.

Cantarel BL, Korf I, Robb SM, Parra G, Ross E, Moore B, Holt C, Alvarado AS, Yandell M. 2008. MAKER: an easy-to-use annotation pipeline designed for emerging model organism genomes. Genome Research 18: 188–196

Chen H, Du L. 2016. Iterative polyketide biosynthesis by modular polyketide synthases in bacteria. Applied Microbiology and Biotechnology 100: 541–557.

Cheng YQ, Tang GL, Shen B. 2003. Type I polyketide synthase requiring a discrete acyltransferase for polyketide biosynthesis. Proceedings of the National Academy of Sciences 100: 3149–3154.

Chooi YH, Krill C, Barrow RA, Chen S, Trengove R, Oliver RP, Solomon PS. 2015. An in planta-expressed polyketide synthase produces (R)-mellein in the wheat pathogen *Parastagonospora nodorum*. Applied and Environmental Microbiology 81:177–186.

Chooi YH, Tang Y. 2012. Navigating the fungal polyketide space: from genes to molecules. The Journal of Organic Chemistry 77: 9933–9953.

Dhillon B, Feau N, Aerts AL, Beauseigle S, Bernier L, Copeland A, Foster A, Gill N, Henrissat B, Herath P, et al. 2015. Horizontal gene transfer and gene dosage drives adaptation to wood colonization in a tree pathogen. Proceedings of the National Academy of Sciences 112: 3451–3456.

Edgar RC. 2004. MUSCLE: multiple sequence alignment with high accuracy and high throughput. Nucleic Acids Research 32: 1792–1797.

Ellison CE, Hall C, Kowbel D, Welch J, Brem RB, Glass NL, Taylor JW. 2011. Population genomics and local adaptation in wild isolates of a model microbial eukaryote. Proceedings of the National Academy of Sciences 108: 2831–2836.

Evans M. 2013. White grain rejection spreading. Ground Cover 102.

Friesen TL, Stuckenbrock EH, Liu Z, Meinhardt S, Ling Hua, Farris JD, Rasmussen JB, Solomon PS, McDonald BA, Oliver RP. 2006. Emergence of a new disease as a result of interspecific virulence gene transfer. Nature Plants 38: 953–956.

Friesen TL, Faris JD, Solomon PS, Oliver RP. 2008. Host-specific toxins: effectors of necrotrophic pathogenicity. Cellular Microbiology 10: 1421–1428.

Goodwin SB, M’Barek SB, Dhillon B, Wittenberg AHJ, Crane CF, Hane JK, Foster AJ, Van der Lee TAJ, Grimwood J, Aerts A, et al. 2011. Finished genome of the fungal wheat pathogen *Mycosphaerella graminicola* reveals dispensome structure, chromosome plasticity, and stealth pathogenesis. PLoS Genetics: dx.doi.org/10.1371/journal.pgen.1002070

Hashimoto M, Nonaka T, Fujii I. 2014. Fungal type III polyketide synthases. Natural Product Reports 31: 1306–1317.

Hertweck C. 2009. The biosynthetic logic of polyketide diversity. Angewandte Chemie International Edition 48: 4688–4716.

Hyde KD, Chomnunti P. Crous PW, Groenewald JZ, Damm U, Ko TWK, Shivas RG, Summerell BA, Tan YP. 2010. A case for reinventory of Australia’s plant pathogens. Personia: Molecular Phylogeny and Evolution of Fungi 25: 50–60.

Islam MS, Haque MS, Islam MM, Emdad EM, Halim A, Hossen QMM, Hossain MZ, Ahmed B, Rahim S, Rahim MS, et al. 2012. Tools to kill: genome of one of the most destructive plant pathogenic fungi *Macrophomina phaseolina*. BMC Genomics 13: 493

Jami F, Slippers B, Wingfield MJ, Gryzenhout M. 2012. Five new species of the Botryosphaeriaceae from Accacia Karroo in South Africa. Cryptogamie Mycologie 33: 245–266

Jami F, Slippers B, Wingfield MJ, Gryzenhout M. 2013. Botryosphaeriaceae species overlap on four unrelated, native South African hosts. Fungal Biology 118: 168–179.

John U, Beszteri B, Derelle E, Van de Peer Y, Read B, Moreau H, Cembella A. 2008. Novel insights into evolution of protistan polyketide synthases through phylogenetic analysis. Protist 159: 21–30.

Keller NP, Turner G, Bennett JW. 2005. Fungal secondary metabolism- from biochemistry to genomics. Nature Reviews Microbiology 3: 937–947

Khaldi N, Collemare J, Lebrun MH, Wolfe KH. 2008. Evidence for horizontal transfer of a secondary metabolite gene cluster between fungi. Genome Biology 9: R18.

Khosla C. 1997. Harnessing the biosynthetic potential of modular polyketide synthases. Chemical Reviews 97: 2577–2590.

Kohli GS, John U, Figueroa RI, Rhodes LL, Harwood DT, Groth M, Bolch CJ, Murray SA. 2015. Polyketide synthesis genes associated with toxin production in two species of *Gambierdiscus* (*Dinophyceae*). BMC Genomics 16:1.

Kohmoto K, Itoh Y, Shimomura N, Kondoh Y, Otani H, Kodama M, Nihsimura S, Nakatsuka S. 1993. Isolation and biological activities of two host-specific toxins from the tangerine pathotype of *Alternaria alternata*. Phytopathology 83: 495–502.

Lohman JR, Ma M, Osipiuk J, Nocek B, Kim Y, Chang C, Cuff M, Mack J, Bigelow L, Li H, et al. 2015. Structural and evolutionary relationships of “AT-less” type 1 Polyketide synthase ketosynthases. Proceedings of the National Academy of Sciences 112: 12693–12698.

Lohse M, Bolger AM, Nagel A, Fernie AR, Lunn JE, Stitt M, Usadel B. 2012. RobiNA: a user-friendly, integrated software solution for RNA-seq-based transcriptomics. Nucleic Acids Research 40: W622–7.

Miyamoto Y, Masunaka A, Tsuge T, Yamamoto M, Ohtani K, Fukumoto T, Gomi K, Peever T, Tada Y, Ichimura K, et al. 2010. *ACTTS3* encoding a polyketide synthase is essential for the biosynthesis of ACT-toxin and pathogenicity in the tangerine pathotype of *Alternaria alternata*. Molecular Plant-Microbe Interactions 23: 406–414.

Morales-Cruz A, Amrine KC, Blanco-Ulate B, Lawrence DP, Travadon R, Rolshausem PE, Baumgartner K, Cantu D. 2015. Distinctive expansion of gene families associated with plant cell wall degradation, secondary metabolism, and nutrient uptake in the genomes of grapevine trunk pathogens. BMC Genomics 16: 1.

Muria-Gonzalez MJ, Chooi YH, Breen S, Solomon PS. 2015. The past, present and future of secondary metabolite research in the Dothideomycetes. Molecular Plant Pathology 16: 92–107

Nakashima T, Ueno T, Funkami H. 1982. Structure elucidation of AK-toxins, host-specific phytotoxic metabolites produced by *Alternaria kikuchiana* Tanaka. Tetrahedron Letters 23: 4469–4472.

Ohm RA, Feau N, Henrissat B, Schoch CL, Horwitz BA, Barry KW, Condon BJ, Copeland AC, Dhillon B, Glaser F, et al. 2012. Diverse lifestyles and strategies of plant pathogenesis encoded in the genomes of eighteen Dothideomycetes fungi. PLoS Pathogens 8: e1003037.

Oliver RP, Solomon PS. 2010. New developments in pathogenicity and virulence of necrotrophs. Current Opinion in Plant Biology 13: 415–419.

Parra G, Bradnam K, Korf I. 2007. CEGMA: a pipeline to accurately annotate core genes in eukaryotic genomes. Bioinformatics 23: 1061–1067.

Platz G. 2011. Wheat and barley disease management in 2011. Yellow spot and head diseases in wheat. Strategies and products for barley leaf rust. GRDC Update Papers, Grains Research and Development Corporation.

Quevillon E, Silventoinen V, Pillai S, Harte N, Mulder N, Apweiler R, Lopez R. 2005. InterProScan: protein domains identifier. Nucleic Acids Research 33: W116–W120.

Schmitt I, Lumbsch HT. 2009. Ancient horizontal gene transfer from bacteria enhances biosynthetic capabilities of fungi. PLoS One 4: e4437.

Shabuer G, Ishida K, Pidot SJ, Roth M, Dahse HM, Hertweck C. 2015. Plant pathogenic anaerobic bacteria use aromatic polyketides to access aerobic territory. Science 350: 670–674.

Shimodaira H, Hasegawa M. 2001. CONSEL: for assessing the confidence of phylogenetic tree selection. Bioinformatics 17: 1246–1247.

Slippers B, Wingfield MJ. 2007. Botryosphaeriaceae as endophytes and latent pathogens of woody plants: diversity, ecology and impact. Fungal Biology Reviews 21: 90–106.

Solovyev, V. 2001. Statistival approaches in eukaryotic gene prediction. Handbook of Statistical Genetics.

Stamatakis A. 2014. RAxML version 8: a tool for phylogenetic analysis and post-analysis of large phylogenies. Bioinformatics btu033.

Stergiopoulos I, Collemare J, Mehrabi R, De Wit PJ. 2013. Phytotoxic secondary metabolites and peptides produced by plant pathogenic Dothideomycete fungi. FEMS Microbiology Reviews 37: 67–93.

Stuckenbrock EH, Christiansen FB, Hansen TT, Dutheil JY, Schierup MH. 2012. Fusion of two divergent fungal individuals led to the recent emergence of a unique widespread pathogen species. Proceedings of the National Academy of Sciences 109: 10954–10959.

Tanaka A, Shiotani H, Yamamoto M, Tsuge T. 1999. Insertional mutagenesis and cloning of the genes required for biosynthesis of the host-specific AK-toxin in the Japanese pear pathotype of *Alternaria alternata*. Molecular Plant-Microbe interactions 12: 691–702.

Thomas G, Jayasena K. 2015. White Grain Disorder of Wheat in Western Australia.

Throckmorton K, Wiemann P, Keller NP. 2015. Evolution of chemical diversity in a group of non-reduced polyketide gene clusters: using phylogenetics to inform the search for novel fungal natural products. Toxins 7: 3572–3607

Thynne E, McDonald MC, Evans M, Wallwork H, Neate S, Solomon PS. 2015. Re-classification of the causal agent of whitegrain disorder on wheat as three separate species of *Eutiarosporella*. Australasian Plant Pathology 44: 527–539.

van der Nest MA, Bihon W, De Vos L, Naidoo K, Roodt D, Rubagotti E, Slippers B, Steenkamp ET, Wilken PM, Wilson A. 2014. Draft genome sequences of *Diplodia sapinea*, *Ceratocystis manginecas*, and *Ceratocystis moniliformis*. IMA Fungus 5: 135–140.

Walton JD. 1996. Host-selective toxins: agents of compatibility. The Plant Cell 8: 1723.

Walton JD. 2006. HC-toxin. Phytochemistry 67: 1406–1413.

Weber T, Blin K, Duddela S, Krug D, Kim HU, Bruccoleri R, Lee SY, Fischback MA, Muller R, Wohlleben W. 2015. antiSMASH 3.0-a comprehensive resource for the genome mining of biosynthetic gene clusters. Nucleic Acids Research 43: W237–W243.

Wenzel SC, Kunze B, Hofle G, Silakowski B, Scharfe M, Blocker H, Muller R. 2005. Structure and biosynthesis of myxochromides S_1-3_ in *Stigmatella aurantiaca*: evidence for an iterative bacterial type 1 polyketide synthase and for module skipping in nonribosomal peptide biosynthesis. ChemBioChem 6: 375–385.

Wildermuth G, Williamson P, Shivas R, McNamara R. 2001. Premature head blight and white grain – a new disease of wheat. Proceedings of the 13^th^ Biennial Australasian Plant Pathology Conference, Queensland Department of Primary Industries.

Yang G, Rose MS, Turgeon BG, Yoder OC. 1996. A polyketide synthase is required for fungal virulence and production of the polyketide T-toxin. The Plant Cell 8: 2139–2150.

Yun CS, Motoyama T, Osada H. 2015. Biosynthesis of the mycotoxin tenuazonic acid by a fungal NRPS-PKS hybrid enzyme. Nature Communications 6.

Zhu G, Laguer MJ, Stejskal F, Millership JJ, Cai X, Keithly JS. 2002. *Cryptosporidium parvum*: the first protist known to encode a putative polyketide synthase. Gene 298: 79–89.

