## Supplementary figures and images for "Acquisition and loss of secondary metabolite clusters shaped the evolutionary path of three recently emerged phytopathogens of wheat"

### Supplementary Materials

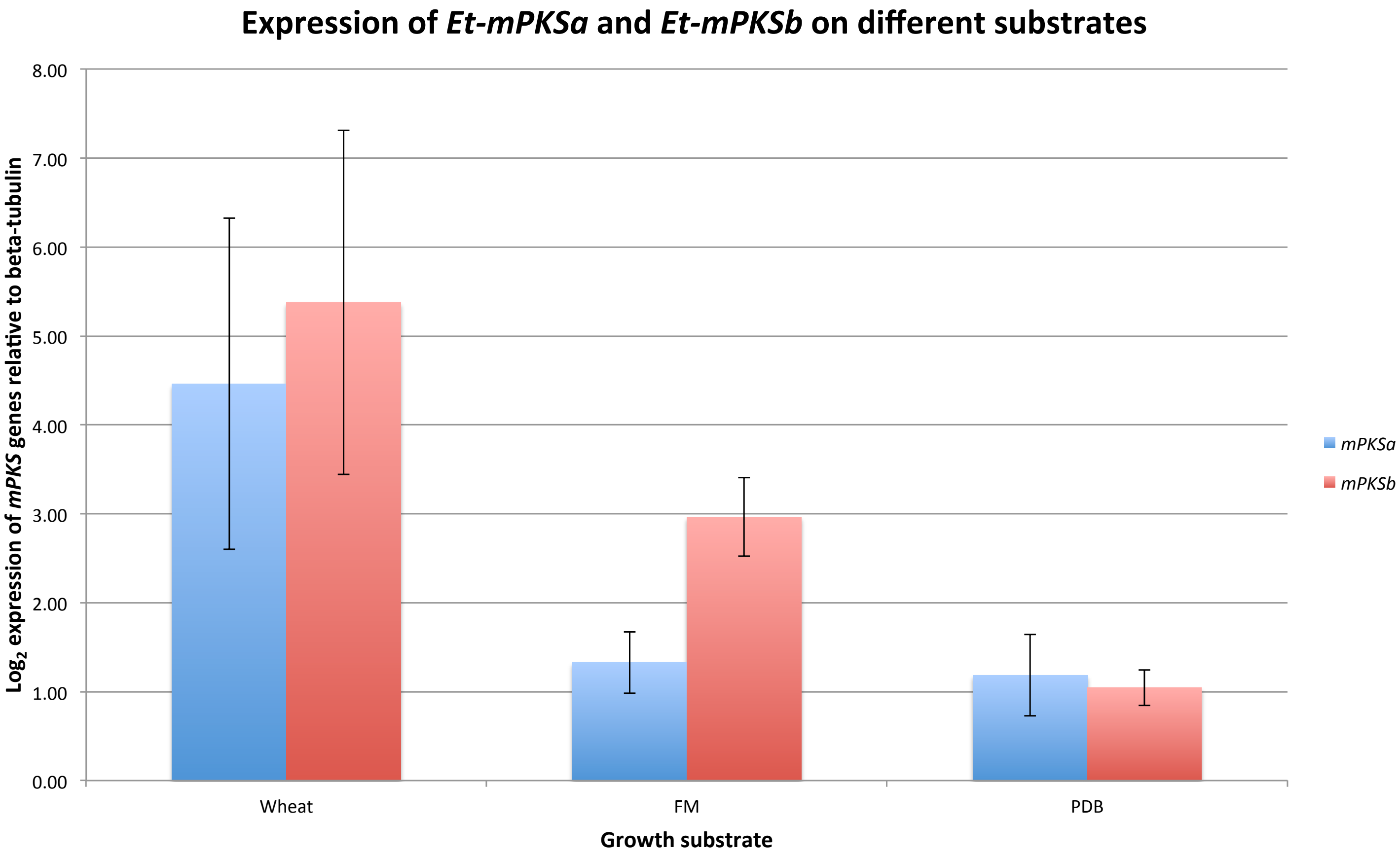

### Supplementary Materials

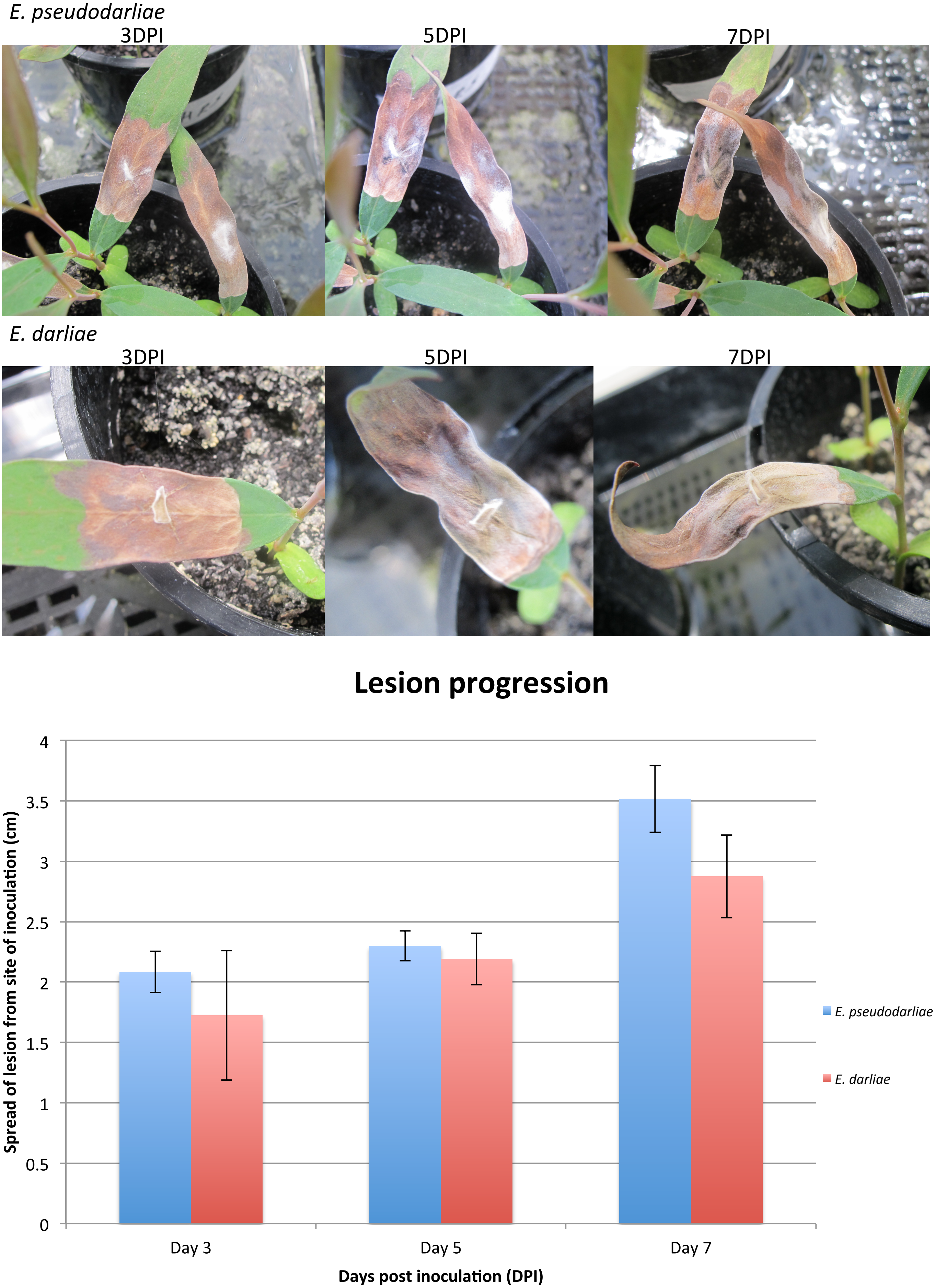

### Supplementary Materials

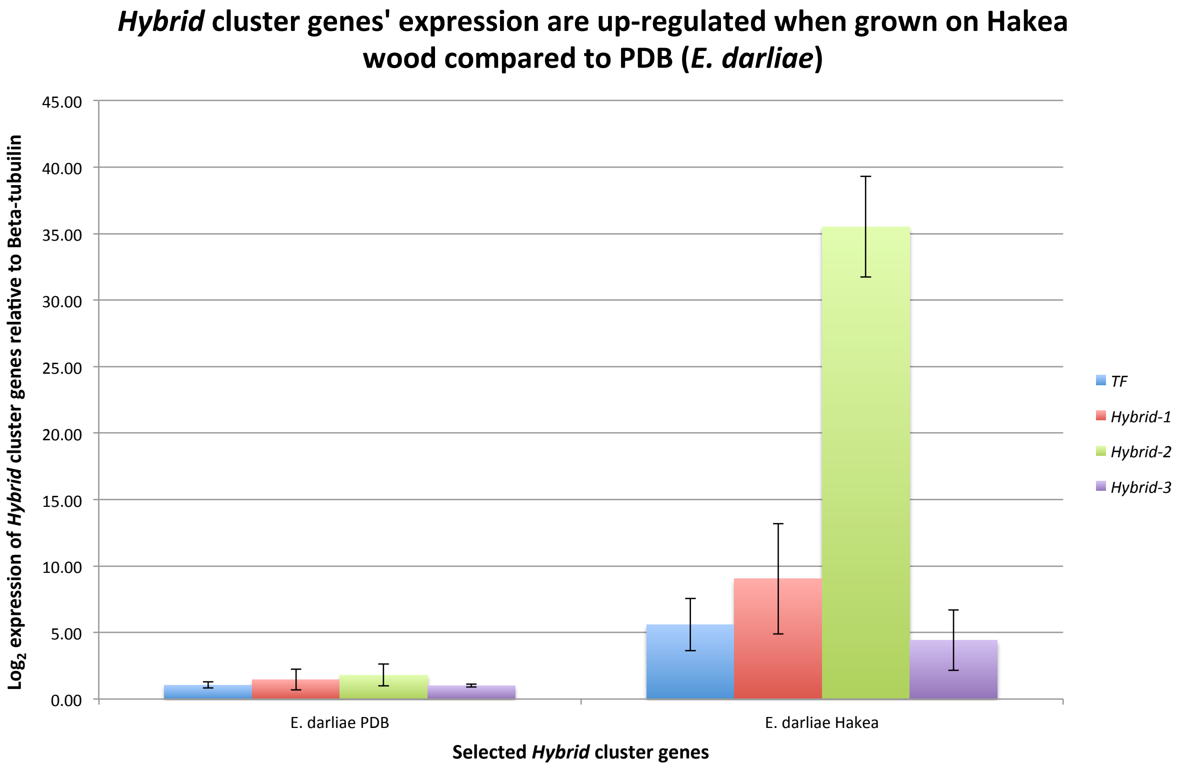

### Supplementary Materials

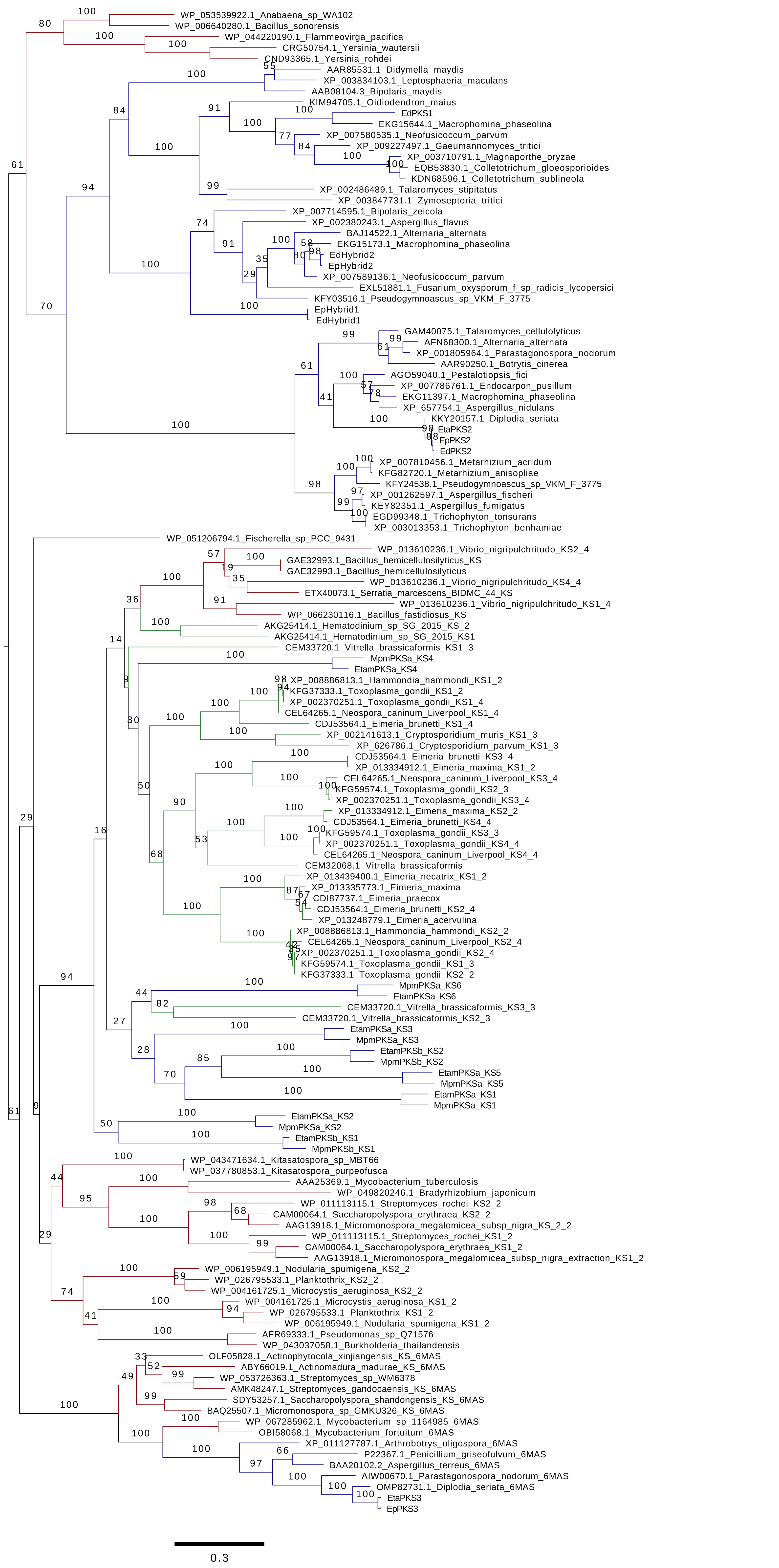

### Supplementary Materials

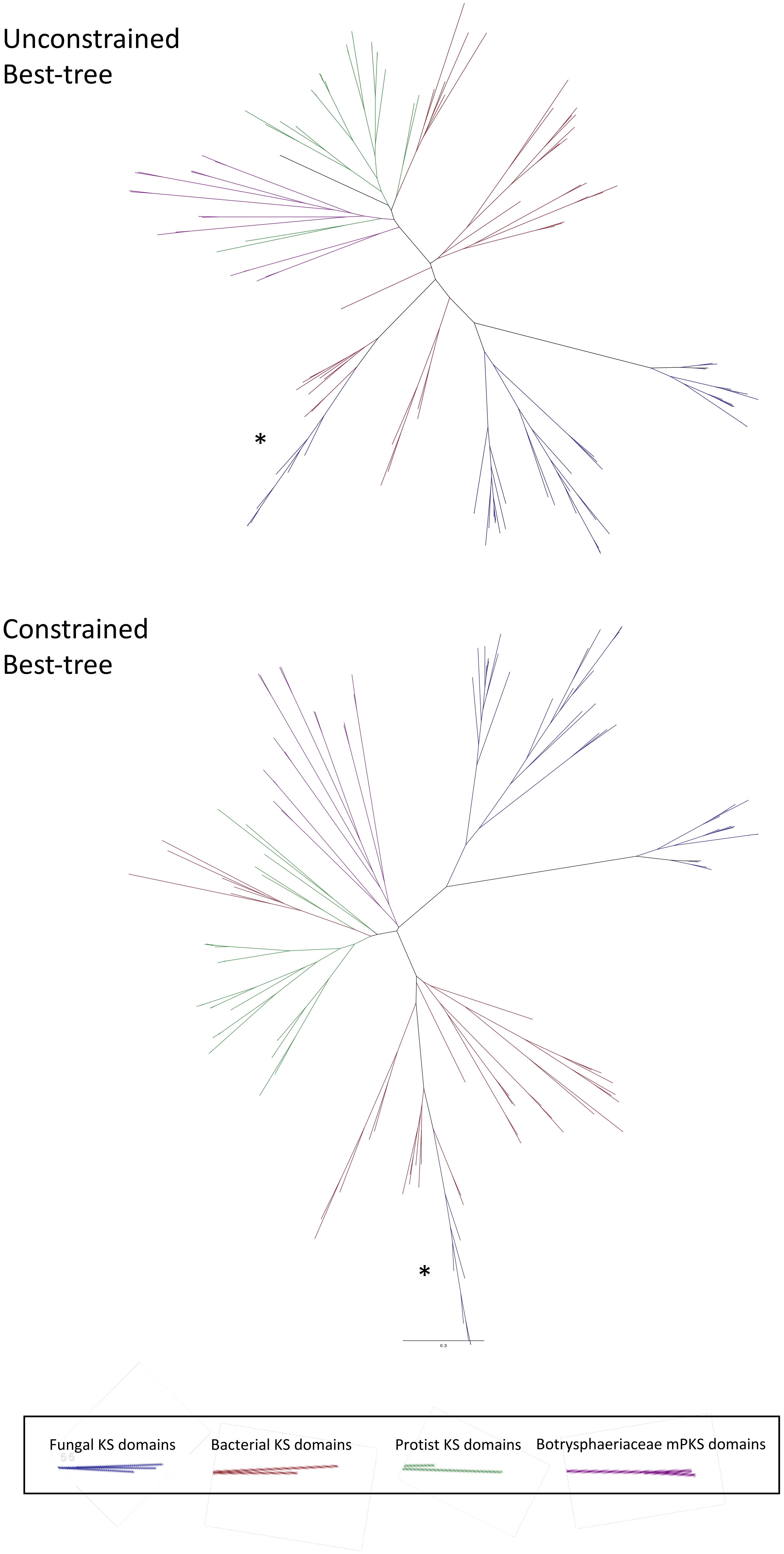

### Supplementary Materials

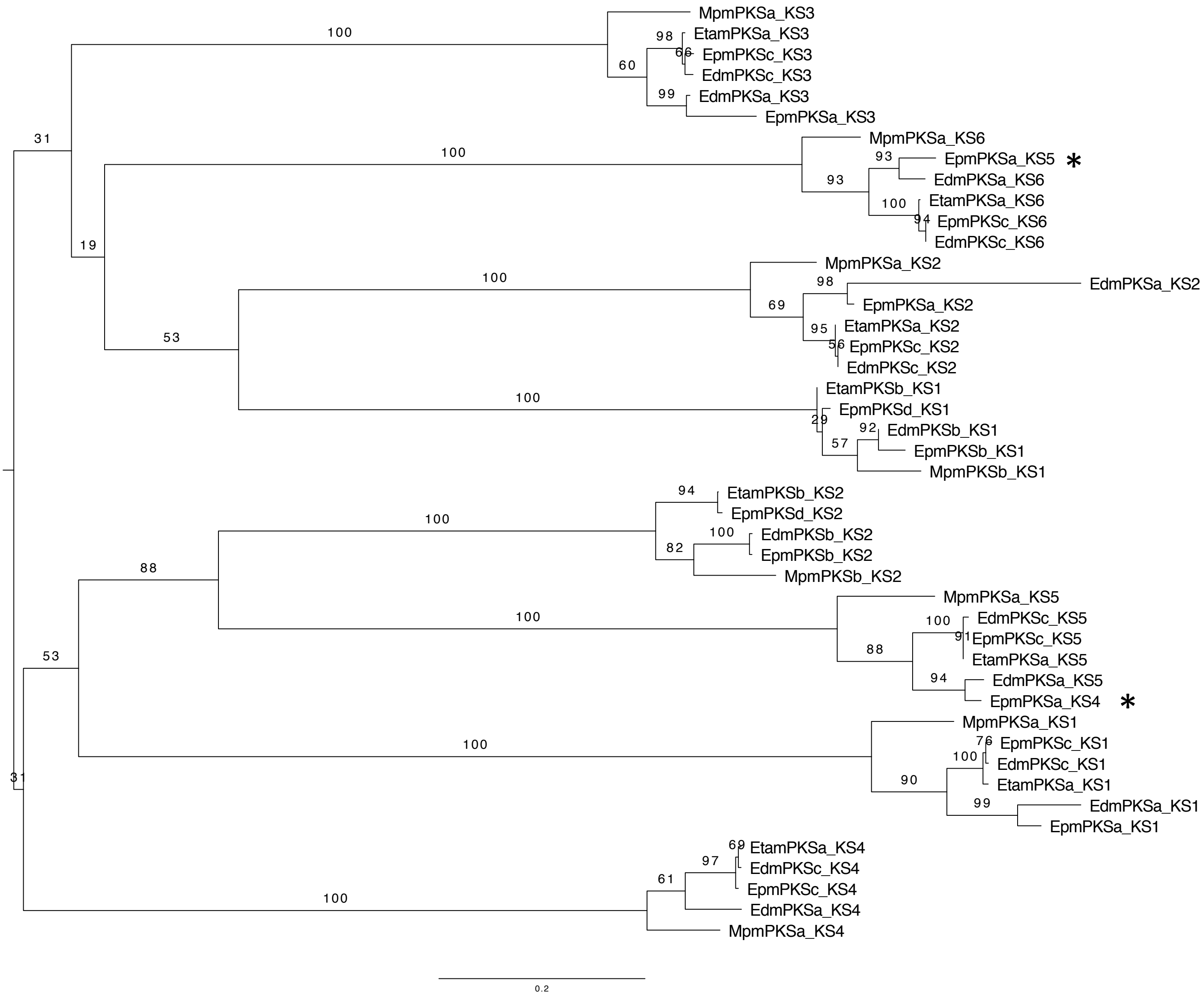

### Supplementary Materials

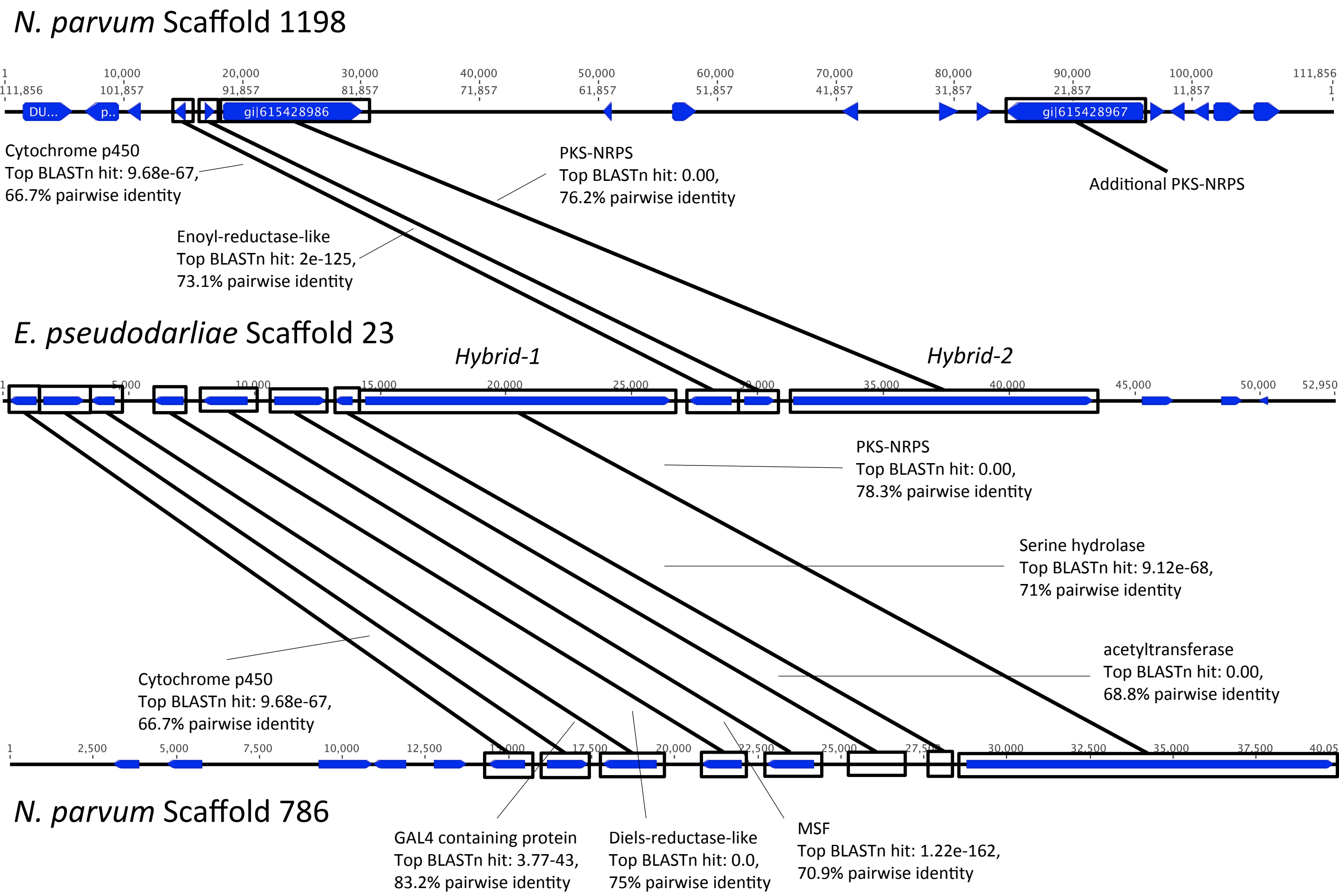

### Supplementary Materials

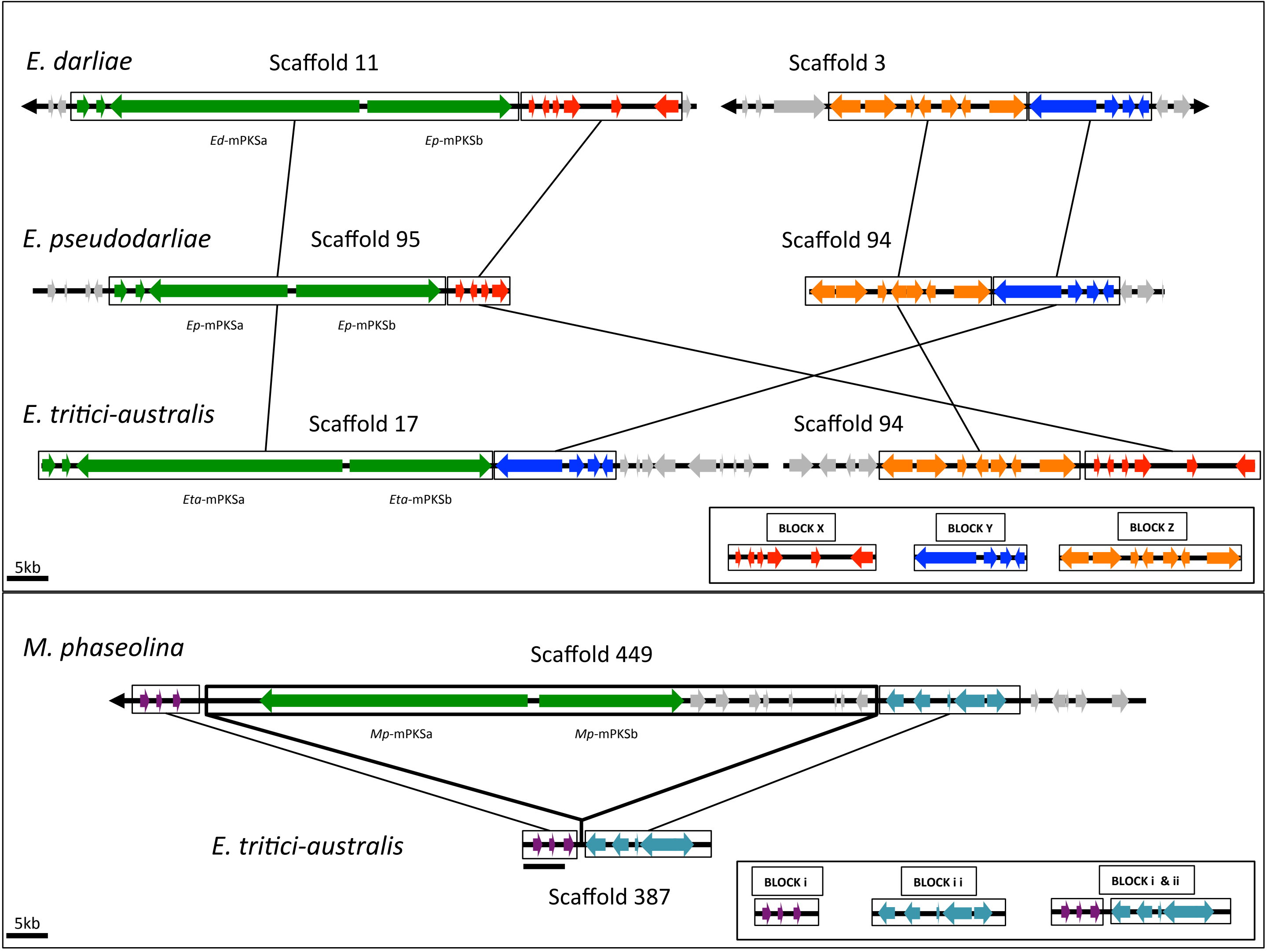

### Supplementary Materials

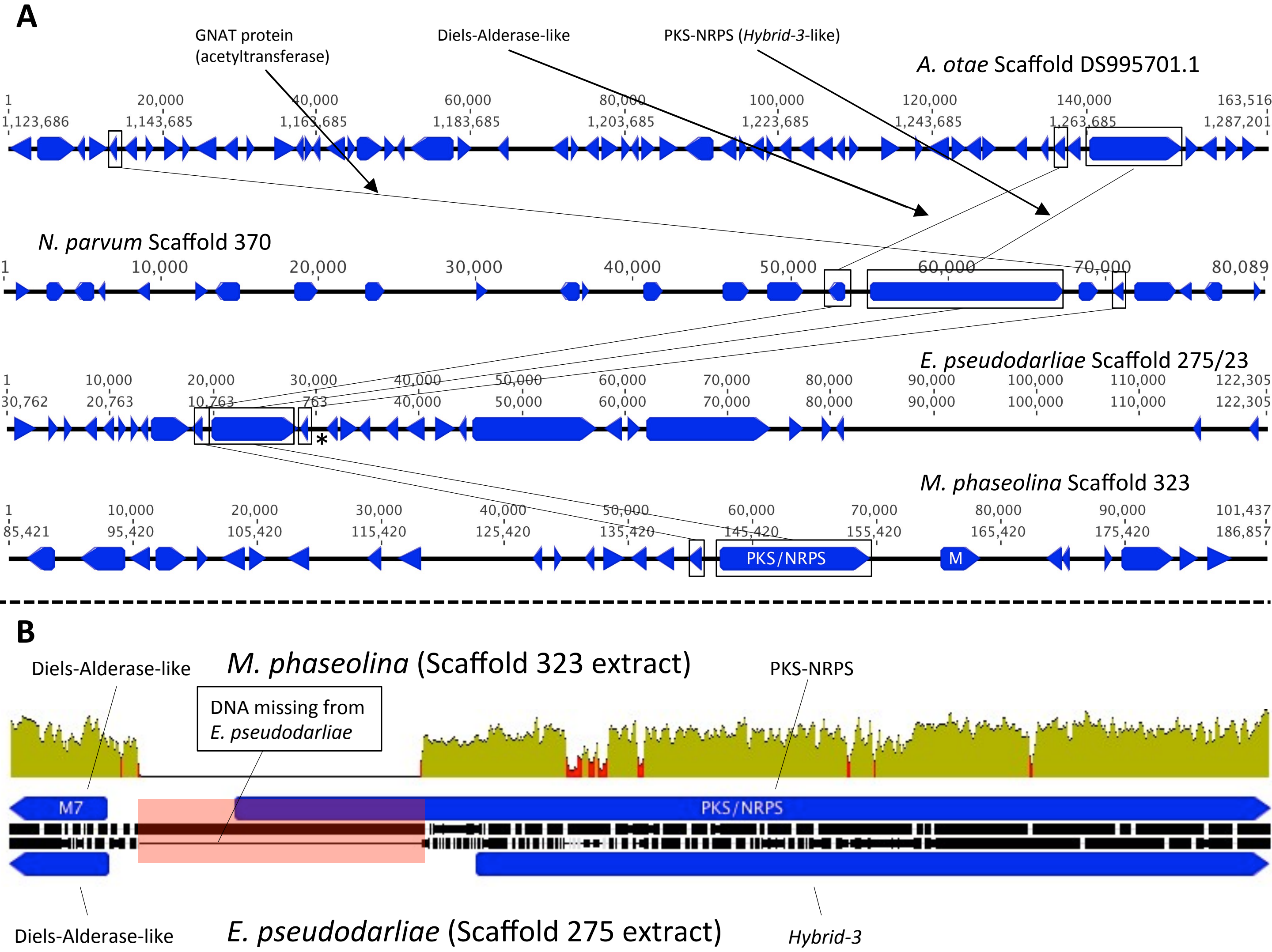

### Supplementary Materials

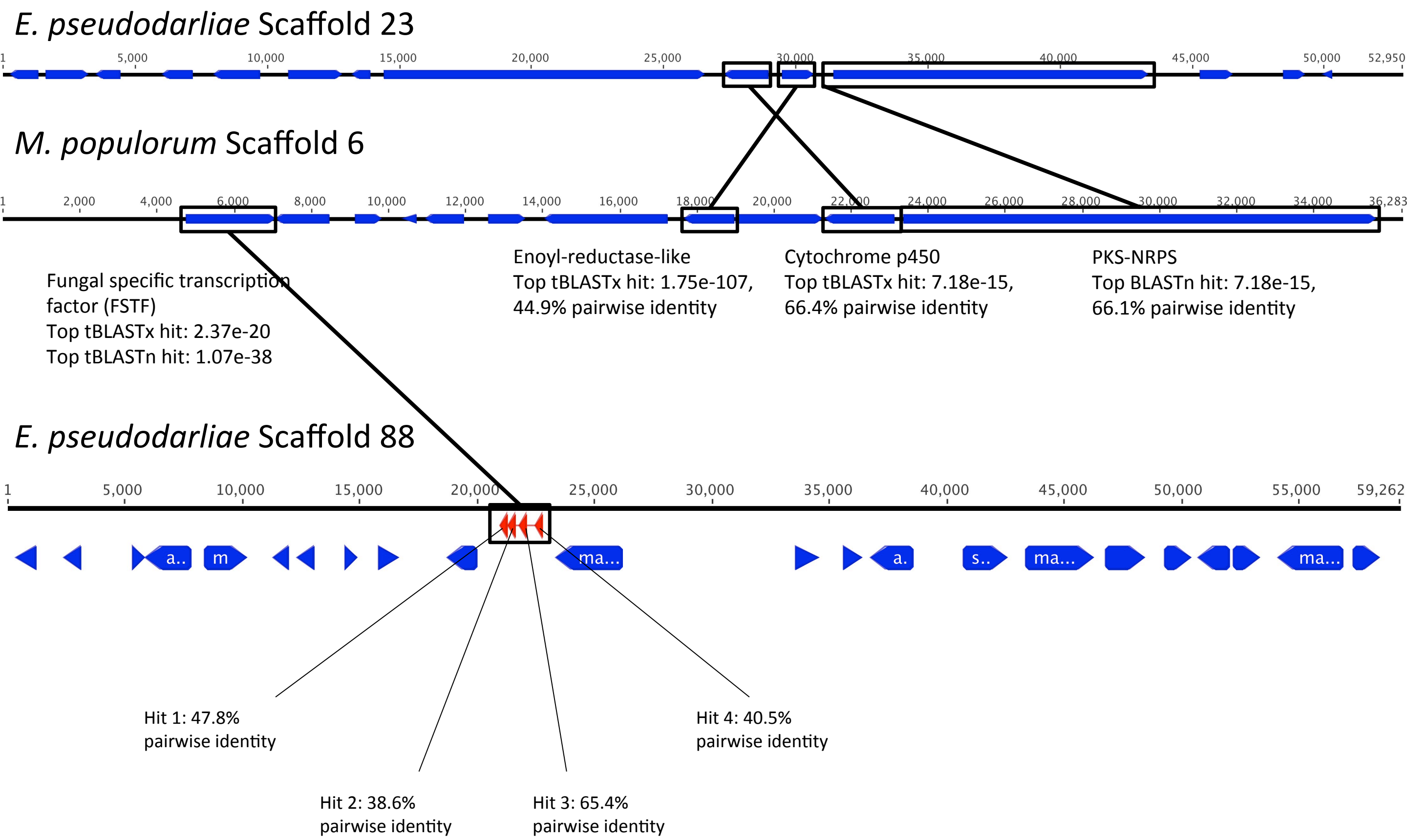

### Supplementary Materials

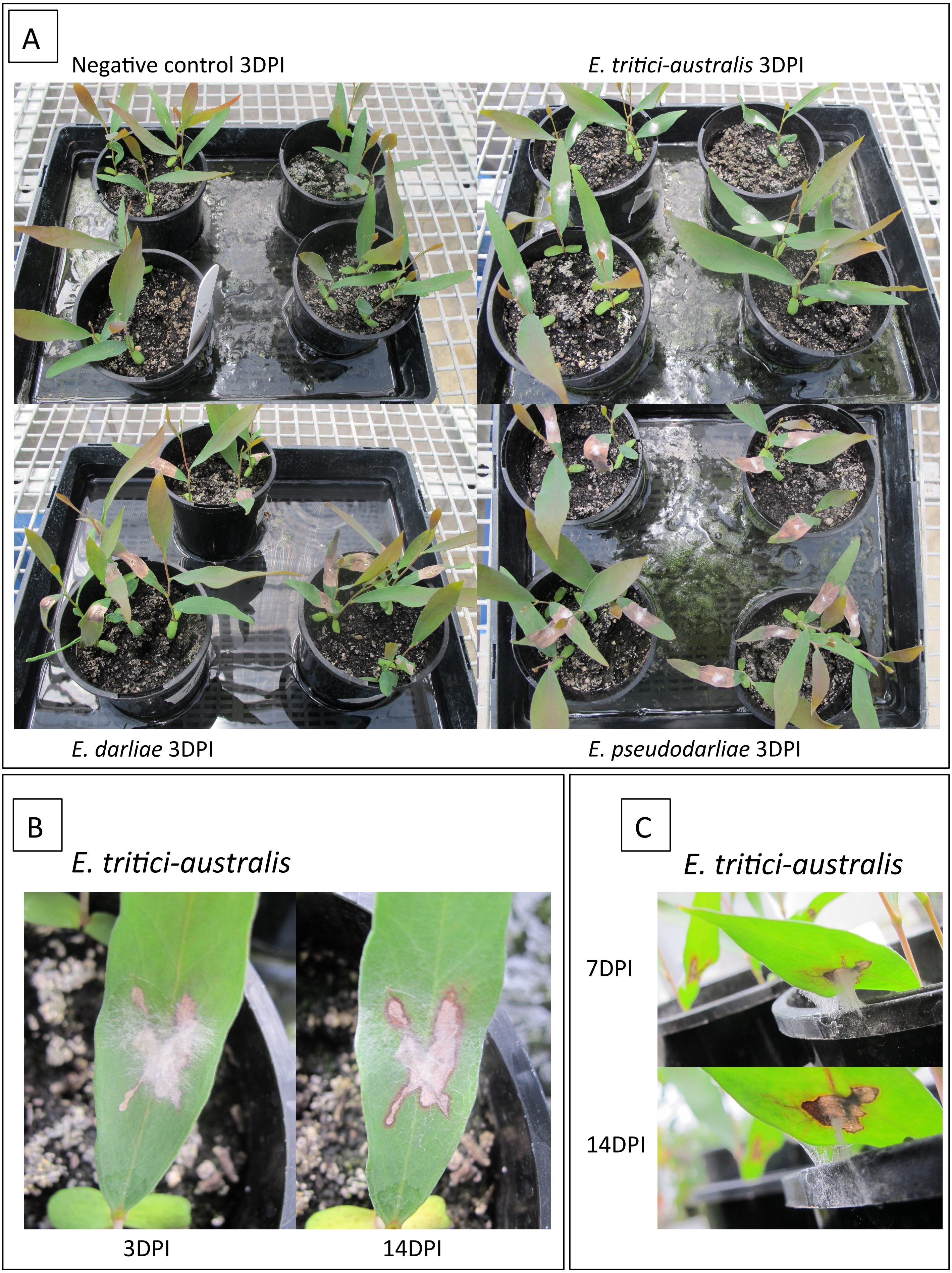
