## Supplementary Materials for "Acquisition and loss of secondary metabolite clusters shaped the evolutionary path of three recently emerged phytopathogens of wheat"

*E. darliae* Isolate 2E2 | *E. darliae* Isolate 2G6 | *E. pseudodarliae* Isolate HR508179 | *E. tritici-australis* Isolate V1-2 | *E. tritici-australis* Isolate V6 | *E. tritici-australis* Isolate V6-1 | *E. tritici-australis* Isolate V18-13 | *E. tritici-australis* Isolate V18-7

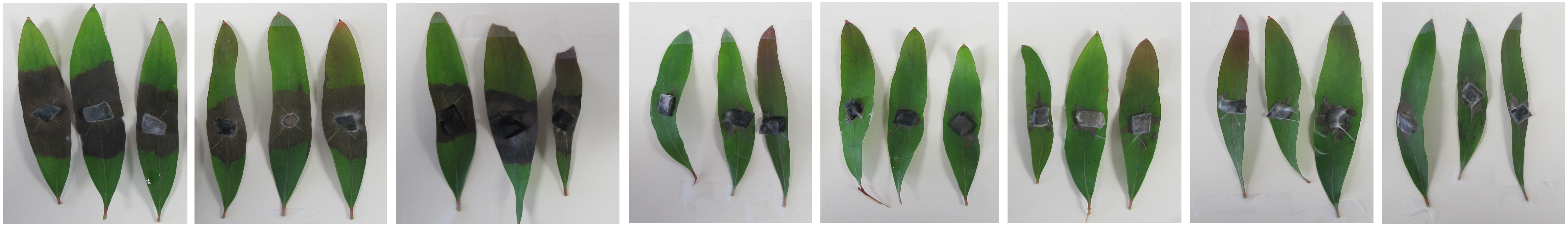
